# Air pollution from biomass burning disrupts early adolescent cortical microarchitecture development

**DOI:** 10.1101/2023.10.21.563430

**Authors:** Katherine L. Bottenhorn, Kirthana Sukumaran, Carlos Cardenas-Iniguez, Rima Habre, Joel Schwartz, Jiu-Chiuan Chen, Megan M. Herting

## Abstract

Exposure to outdoor particulate matter (PM_2.5_) represents a ubiquitous threat to human health, and particularly the neurotoxic effects of PM_2.5_ from multiple sources may disrupt neurodevelopment. Studies addressing neurodevelopmental implications of PM exposure have been limited by small, geographically limited samples and largely focus either on macroscale cortical morphology or postmortem histological staining and total PM mass. Here, we leverage residentially assigned exposure to six, data-driven sources of PM_2.5_ and neuroimaging data from the longitudinal Adolescent Brain Cognitive Development Study (ABCD Study®), collected from 21 different recruitment sites across the United States. To contribute an interpretable and actionable assessment of the role of air pollution in the developing brain, we identified alterations in cortical microstructure development associated with exposure to specific sources of PM_2.5_ using multivariate, partial least squares analyses. Specifically, average annual exposure (i.e., at ages 8-10 years) to PM_2.5_ from biomass burning was related to differences in neurite development across the cortex between 9 and 13 years of age.

**Graphical Abstract:** 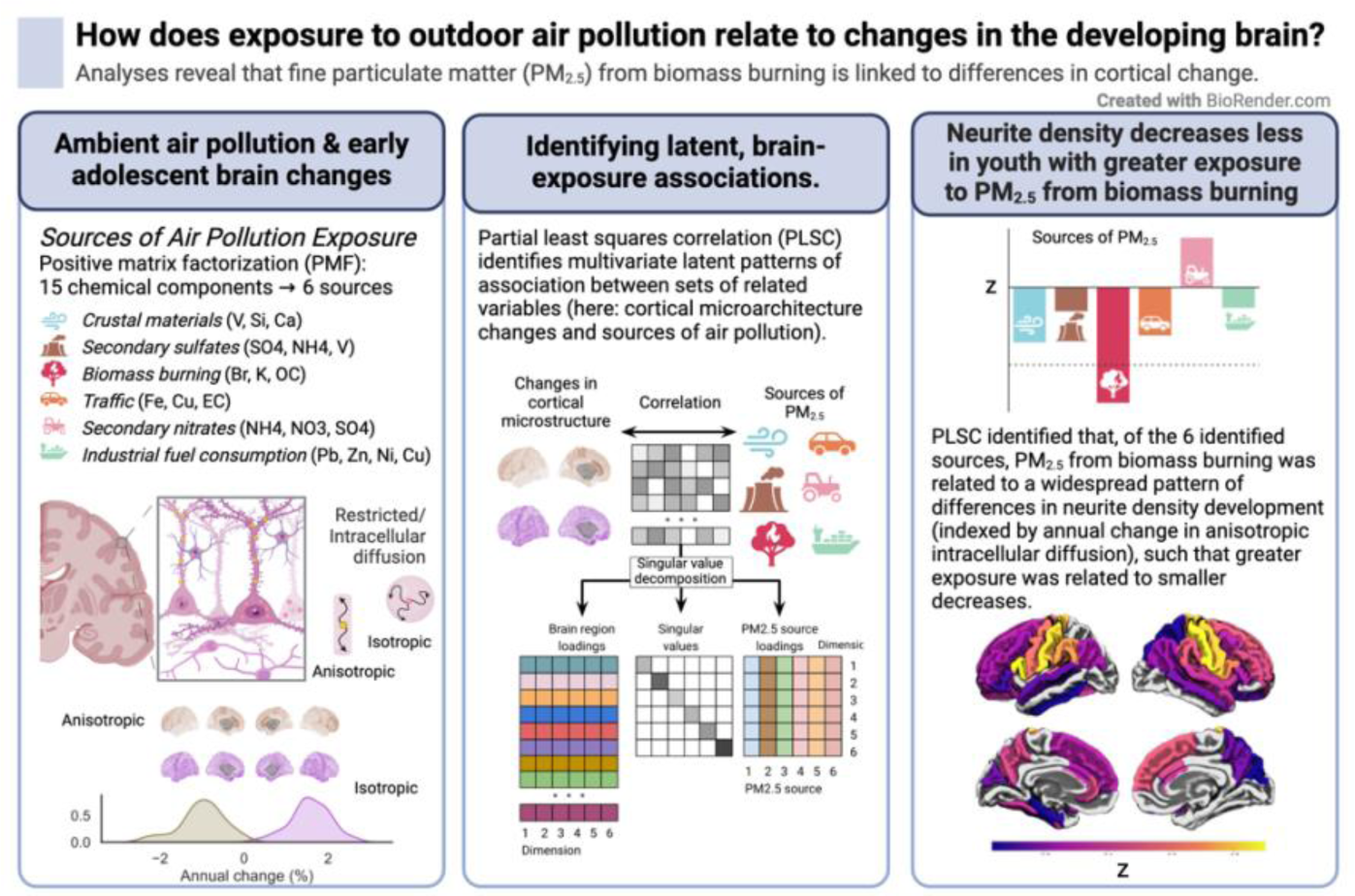

## 1. Introduction

Outdoor air pollution and, in particular, exposure to fine particulate matter with aerodynamic diameter less than 2.5µm (i.e., PM_2.5_), is among the greatest threats to human health due to its ubiquity and widespread effects (Cardenas-Iniguez et al., 2022; Costa et al., 2017; Genc et al., 2012). These effects are exacerbated in children (Brumberg et al., 2021), who have greater respiratory rates, are usually more physically active, and spend more time outdoors than adults, incurring greater dose and greater effect (Bateson and Schwartz, 2007). In addition to its neurotoxic effects, PM_2.5_ exposure has been associated with a number of poor behavioral outcomes, including increased depression and suicide risk (Braithwaite et al., 2019; Heo et al., 2021; Liu et al., 2021), anxiety and psychosis (Newbury et al., 2021), and poor performance on cognitive assessments (Allen et al., 2017b; Clifford et al., 2016; Guxens et al., 2018, 2014), albeit findings are mixed (Essers et al., 2023; Kusters et al., 2022). Together, the risk of PM_2.5_ exposure and breadth of potential neurodevelopmental consequences highlight the importance of leveraging novel neuroimaging methods to elucidate neurotoxicant effects of air pollution on the developing human brain. One challenge in this line of research is that PM_2.5_ is a complex mixture of chemical components (e.g., metals, nitrates, sulfates, carbons) arising from sources including anthropogenic human activities (e.g., traffic, industrial fuel burning), natural and meteorological events (e.g., windblown dust, wildfires). The sources and composition of PM_2.5_ vary geographically (Snider et al., 2016) and can have different effects on human health (C. Chen et al., 2021; Chung et al., 2015; Holguin, 2008; Kazemiparkouhi et al., 2022; Sarnat et al., 2008). Untangling the effects of different sources of PM_2.5_ (e.g., via data-driven source apportionment) is important for understanding the effects of outdoor air pollution exposure on human health and neurodevelopment. Source apportionment analyses can help provide insight as to the origins of outdoor PM_2.5_, which may be helpful in designing mitigation strategies and policies to improve air quality. Furthermore, it could inform future experimental designs to study the mechanisms of “real world” exposure settings (Godleski et al., 2011), while providing more precise estimates of the global burden of disease associated with PM_2.5_ (McDuffie et al., 2021). Overall, mapping neurotoxic effects of PM_2.5_ exposure to sources may facilitate a clearer picture of its neurodevelopmental impacts and provide actionable insights for researchers, parents, and policymakers.

In addition to increased exposure, children are likely more susceptible to longer-term effects of PM_2.5_ exposure when it affects their ongoing development. Brain development, in particular, follows a protracted course and continues into the third decade of life (Herting et al., 2018, 2017; Herting and Sowell, 2017). The protracted course of brain maturation presents a large window of opportunity for air pollution and PM_2.5_ more specifically to impact neurodevelopment (Brockmeyer and d’Angiulli, 2016; Herting et al., 2019). Notably, neurotoxic effects of PM_2.5_ are thought to include cellular processes that are ongoing throughout childhood and early adolescence, including apoptosis (Wang et al., 2021), changes in dendritic spine density and arborization (Allen et al., 2017a; Fonken et al., 2011), neurogenesis (Woodward et al., 2018), and microglial functioning (Allen et al., 2017a; Cardenas-Iniguez et al., 2022; Costa et al., 2017; Genc et al., 2012). Thus, it is crucial to consider not only individual differences in brain structure and function, but how developmental *changes* relate to different sources of PM_2.5_ exposure, to better understand its neurodevelopmental effects. In this realm, most research to date has linked PM_2.5_ exposure to cross-sectional differences in cortical morphology, white matter microarchitecture, and subcortical microarchitecture. Urban air pollution, which includes PM_2.5_, exposure has been associated with increased white matter hyperintensities (Calderón-Garcidueñas et al., 2012, 2008), neurovascular dysfunction and ultrafine particle deposition in the brain (Calderón-Garcidueñas et al., 2016, 2008), metabolic alterations (Calderón-Garcidueñas et al., 2015), and altered white matter microarchitecture in children (Binter et al., 2022; Burnor et al., 2021; Huuskonen et al., 2021). Thinner cortex and smaller subcortical volumes have been associated with greater exposure to ambient PM_2.5_, as well (de Prado Bert et al., 2018; Herting et al., 2019; Lubczyńska et al., 2021; Peterson et al., 2022). While exposure to PM_2.5_ has documented neurotoxic effects and there is substantial evidence of differential health effects between sources of PM_2.5_, little research has considered source-specific impacts on the brain. Thus, this work addresses two key gaps in our understanding of the neurodevelopmental implications of PM2.5 exposure first, regarding impacts of exposure on cortical microarchitecture changes and, second, regarding the implications of exposure to different sources of PM_2.5_ for brain development.

Fine PM, or PM_2.5_, can cross the blood brain barrier (Kang et al., 2021) and, as previously mentioned, animal models have shown that it can induce various cellular processes including neuroinflammation, apoptosis, altered dendritic spine density and arborization, and neurogenesis (Cory-Slechta et al., 2023; Ferreira et al., 2022; Liu et al., 2023). Thus, it is feasible that exposure during development may not only impact morphology, but microstructural properties of cortical gray matter tissue. Given our understanding of the cellular impacts of PM in the brain, assessing gray matter microarchitecture could provide important neurobiological insight into *in vivo* impacts of PM exposure in humans. Two studies to date have linked PM_2.5_ exposure to subcortical gray matter microarchitecture using diffusion-weighted magnetic resonance imaging (dMRI) approaches, revealing differences in the thalamus, brainstem, nucleus accumbens, and caudate nucleus (Peterson et al., 2022; Sukumaran et al., 2023). However, little is known about its effects on human *cortical* gray matter microarchitecture. Recent advances in dMRI facilitate multi-compartment, biophysical models that can provide more detailed estimates of cortical gray matter microarchitecture (Martinez-Heras et al., 2021). Restriction spectrum imaging (RSI) takes advantage of multi-shell diffusion MRI data to separately model diffusion in intracellular and extracellular compartments (White et al., 2012). Within intracellular diffusion are anisotropic and isotropic components, that separately estimate cylindrical diffusion, along a primary axis or axes, and spherical diffusion, respectively. Further, histological validation has shown that intracellular RSI measures can provide estimates of cortical neurite (i.e., axon and dendrite) architecture and cellularity (White et al., 2012), and may be sensitive to neurobiological processes underlying development during late childhood and early adolescence (e.g., synaptic pruning, apoptosis) (Palmer et al., 2022). During early adolescence, neurite density, as indexed by anisotropic intracellular diffusion, largely decreases across the cortex, while cellularity, as indexed by isotropic intracellular diffusion, largely increases (Bottenhorn et al., 2023). As the protracted nature of cortical development creates the largest window of vulnerability to lasting neurotoxic effects of air pollution across the brain, such a perspective could provide important information regarding the translation of childhood exposure into lasting damage to brain and mental health.

Here, we assess source-specific impacts of average annual PM_2.5_ exposure on cortical microarchitecture changes in children between ages 9-10 years and two-years later from the longitudinal, nationwide Adolescent Brain Cognitive Development Study (ABCD Study®). The ABCD Study enrolled 11,881 children from 21 data collection sites across the United States and includes estimates of 15 chemical components of PM_2.5_ linked to their residential addresses, with 50m^2^ resolution in urban areas and 1km^2^ resolution elsewhere. From these components, we used positive matrix factorization to identify six sources of PM_2.5_ (Sukumaran et al,. in preparation). We chose to assess how one-year average exposure to PM_2.5_ sources during childhood (i.e., between ages 8 and 10 years) were related to changes in cortical microarchitecture between ages 9 and 13 years. Cortical microstructure was estimated from both anisotropic and isotropic intracellular diffusion, reflecting neurite density and cellularity, respectively (Carper et al., 2017; Conley et al., 2021; Newman et al., 2023; Palmer et al., 2022; Rapuano et al., 2020), as hypothesized neurotoxic effects of PM_2.5_ likely include disruption to ongoing synaptic refinement during adolescence and altered cellularity due to neuron death and/or reactive gliosis. Using Partial Least Squares Correlation (PLSC), we jointly modeled latent dimensions of association between source-specific exposure to PM_2.5_ and changes in cortical microarchitecture. By investigating source-specific effects on these aspects of early adolescent neurodevelopment, this work aims to illuminate a more mechanistic and policy-relevant understanding of the developmental neurotoxicity of outdoor air pollution. We expect one or more sources of ambient air pollution to be associated with individual differences in cortical microarchitecture changes and that latent associations between sources of air pollution and cortical microstructure will likely include regions changing the most during this developmental period but vary across brain regions.

## 2. Material and methods

### 2.1 Participants

Longitudinal data from the ongoing ABCD Study® were obtained from the annual 4.0 and 5.0 data releases (http://dx.doi.org/10.15154/1523041; https://dx.doi.org/10.15154/8873-zj65; Supplementary Table 0). The ABCD Study enrolled over 11,800 children 9 to 10 years of age in a 10-year longitudinal study (Garavan et al., 2018). Participants were recruited at 21 study sites across the United States, largely from elementary schools (private, public, and charter schools) in a sampling design that aimed to represent the nationwide sociodemographic diversity (Casey et al., 2018). All experimental and consent procedures were approved by the institutional review board and human research protections programs at the University of California San Diego. Each participant provided written assent to participate in the study and their legal guardian provided written agreement to participate. Here, we use a subset of data from the ABCD Study, including magnetic resonance imaging (MRI), in addition to measures of participants’ sex at birth, socio-demographics, and air pollution exposure. Neuroimaging data include assessments from two time points: baseline enrollment and year 2 follow-up. Sociodemographic and exposure data include assessments from the baseline time point. Exclusion criteria for the ABCD study included lack of English proficiency, severe sensory, neurological, medical or intellectual limitations, and inability to complete an MRI scan. For more information, see Garavan et al. (2018) and Volkow et al. (Volkow et al., 2018).

For this study, we further excluded participants whose 2-year follow-up visit occurred after shutdowns associated with the coronavirus pandemic (i.e., March 2020). Shutdowns in response to the COVID-19 pandemic changed the air quality across the United States (Bekbulat et al., 2021; K. L. Chen et al., 2021) while also disrupting the lives of youth. Imaging data was excluded if any of the following criteria were met: it had incidental neurological findings evident in their scans, failed the ABCD imaging or FreeSurfer quality control procedures (Hagler et al., 2019), and/or was missing diffusion-weighted imaging data from either baseline or 2-year follow-up time point, or greater than 2.0 mm of head motion (i.e., framewise displacement) during a diffusion-weighted scan.

### 2.2 Demographic Data, Confounders, & Precision Variables

In terms of age, sex, and household size, the ABCD Study cohort closely matches the distribution of 9- and 10-year-olds in the American Community Survey (ACS), a large probability sample survey of U.S. households conducted annually by the U.S. Bureau of Census (Compton et al., 2019). The racial breakdown matches closely, too, although children of Asian, American Indian/Alaska Native and Native Hawaiian/Pacific Islander ancestry are under-represented in ABCD (Heeringa and Berglund, 2020). Confounders and precision variables to be adjusted for in our analyses were identified using a directed acyclic graph (DAG; Supplementary Figure 1) (Shrier and Platt, 2008; Textor et al., 2016) constructed with theoretical knowledge of variables associated with brain development and air pollution exposure. From this DAG, a minimally sufficient set of covariates were identified, which represent the smallest group of variables that can account for all paths by which the entire variable set may impact air pollution exposure based on the primary residential address and cortical microarchitecture development. Variables included the child’s age at baseline data collection (months), sex assigned at birth (*male* or *female*), caregiver-reported race/ethnicity (*White, Black, Hispanic, Asian,* or *Other*), and handedness (*right, left,* or *mixed*; (Oldfield, 1971; Veale, 2014)), average daily screen time (hours; (Paulich et al., 2021; Paulus et al., 2019)), physical activity (number of days child was physically active in the week prior; (Barch et al., 2018)), combined household income in USD (*>100K, 50-100K, <50K,* or *Don’t Know/Refuse to Answer*), caregiver-reported perceived neighborhood safety based on a survey modified from PhenX (NSC) (Echeverria et al., 2004; Mujahid et al., 2007), and the following information about the child’s primary residence: the population density (“Center for International Earth Science Information Network,” n.d.; Fan et al., 2021), urbanicity (*US Census Track Urban Classification*), distance to major roadways (meters; (Fan et al., 2021)), and nighttime noise (i.e., average noise between the hours of 10:00 PM and 7:00 AM in decibels; (Mennitt et al., 2014)). The manufacturer of the MRI on which each participant’s data were collected and the average head motion (framewise displacement, mm) throughout diffusion scans at each time point were included in the minimally sufficient set, as well.

### 2.3 Neuroimaging Data

#### 2.3.1 MRI: Acquisition, Processing, and Quality Control

A harmonized data protocol was utilized across sites with either a Siemens, Phillips, or GE 3T MRI scanner (Casey et al., 2018). Motion compliance training, as well as real-time, prospective motion correction was used to reduce motion distortion (Casey et al., 2018). T1w images were acquired using a magnetization-prepared rapid acquisition gradient echo (MPRAGE) sequence and T2w images were obtained with a fast spin echo sequence with variable flip angle. Both consist of 176 slices with 1mm^3^ isotropic resolution. The DWI acquisition included a voxel size of 1.7 mm isotropic and implements multiband EPI with slice acceleration factor 3. Each DWI acquisition included a fieldmap scan for B0 distortion correction. ABCD employs a multi-shell diffusion acquisition protocol that includes 7 b=0 frames as well as 96 total diffusion directions at 4 b-values (6 with b = 500 s/mm^2^, 15 with b = 1000 s/mm^2^, 15 with b = 2000 s/mm^2^, and 60 with b = 3000 s/mm^2^). For more details on the scanning protocol, please see Casey et al. (2018). All images underwent distortion correction, bias field correction, motion correction, and manual and automated quality control per the steps detailed by Hagler and colleagues (2019). Only images without clinically significant incidental findings (*mrif_score* = 1 or 2) that passed all ABCD Study quality-control parameters were included in analysis (*imgincl_dmri_include* = 1). For a more detailed description of diffusion data processing, please refer to recent work by Gorham & Barch and Hagler et al. (Gorham and Barch, 2020; Hagler et al., 2019).

#### 2.3.2 Restriction Spectrum Imaging (RSI)

Restriction spectrum imaging (RSI) is an advanced modeling technique that utilizes all 96 directions collected as part of the ABCD Study’s multi-shell, high angular resolution imaging (HARDI) acquisition protocol. RSI provides detailed information regarding both the extracellular and intracellular compartments of white matter within the brain (White et al., 2014, 2013, 2012). RSI model outputs include normalized measures of intracellular (i.e., restricted), extracellular (i.e., hindered), and free water movement, all of which are on a unitless scale of 0 to 1. Restricted normalized isotropic signal fraction (RNI), or *isotropic intracellular diffusion,* measures directionless water movement at short distances such that a higher RNI could indicate an increase in number of neuronal cell bodies (i.e., neurogenesis) and/or support cells or swelling of support cells, such as activated astrocytes or microglia as in neuroinflammation (Palmer et al., 2022). Restricted normalized directional signal fraction (RND), or *anisotropic intracellular diffusion,* measures directional water movement at short distances and likely indicates intra-neurite (i.e., axons and dendrites) diffusion, with higher RND indicating more myelination, axonal packing, and/or dendritic arborization (Palmer et al., 2022).. Mean RSI measures are calculated for all cortical ROIs from the Desikan-Killiany atlas (Desikan et al., 2006) to provide estimates of cortical microarchitecture across the brain.

#### 2.3.3 Annualized Percent Change in Cortical Microarchitecture

As previously published in greater detail, annualized percent change between baseline and 2-year follow up data collection time points was calculated to capture intra-individual changes in cortical microarchitecture (Bottenhorn et al., 2023).

*Annualized percent change* = [((*measure_T2_* − *measure_T1_*) ÷ (*measure_T2_* + *measure_T1_*) / 2) × 100] ÷ (*age_T2_* − *age_T1_*), where Tt_1_ = time point 1 (i.e., baseline data collection visit, between ages 9-10 years) and Tt_2_ = time point 2 (i.e., 2-year follow-up visit, between ages 11-13 years).

### 2.4 Particulate Matter Components and Source Groups

The estimates of outdoor PM_2.5_ exposure represent averages over the initial year of ABCD Study data collection (i.e., 2016) and were compiled and linked based on participants’ residential addresses by the ABCD Study’s LED workgroup (Fan et al., 2021). At participants’ initial study visit, between October 2016 and October 2018, each participant’s caregiver reported their primary residential addresses, which were later geocoded to estimate environmental exposures, among other variables. Daily outdoor PM_2.5_ exposure was estimated in µg/m^3^ at 1km^2^ resolution via novel, hybrid, machine-learning based, spatiotemporal models that leverage satellite-based aerosol optical depth observations, along with land-use regression and chemical transport model outputs (Di et al., 2019). These ensemble models included neural networks, random forest, and gradient boosting algorithms and were cross-validated with EPA measurements across the U.S., demonstrating excellent performance (spatial R2 = 0.89, spatial RMSE = 1.26µg/m3, temporal R2 = 0.85) (Di, Amini, Shi, Kloog, Silvern, Kelly, Sabath, Choirat, Koutrakis, & Lyapustin, 2019). These daily exposures were, then, averaged over the first year of baseline data collection (2016) to estimate average annual exposure. A similar method was used to estimate 15 separate PM_2.5_ components across the U.S. at a 50m^2^ spatial resolution: elemental carbon (EC), organic carbon (OC), silicon (Si), potassium (K), calcium (Ca), bromine (Br), nitrate (NO_3_^−^), ammonium (NH_4_^+^), sulfate (SO_4_^2−^), vanadium (V), iron (Fe), nickel (Ni), copper (Cu), zinc (Zn), and lead (Pb). Distributions of the mass concentrations for each component are available in Supplementary Figure 3, while pairwise Spearman correlations between participant exposures to each of these components are available in Supplementary Figure 4. In estimating each component, 166 predictors were used, including temporal and geographical information, observations from satellite (e.g., aerosol optical depth, nighttime lights, vegetation, water index) and meteorological data (e.g., humidity, temperature, wind), emission sources or surrogates thereof (e.g., distance to power plants and highways, traffic counts) (Amini et al., 2022; Jin et al., 2022). For model performance per component, see Supplementary Table 1. As part of a larger scope of ongoing work, we performed source apportionment on these annual estimates of PM_2.5_ components via positive matrix factorization (PMF) using the PMF tool developed and released by United States Environmental Protection Agency (EPA, v5.0) (Sukumaran et al., Under Review). Briefly, PMF decomposes the 15 PM_2.5_ components into a predetermined number of sources (or factors), while constraining both the factors and components’ contributions to each factor to non-significantly negative (i.e. positive) values. When repeated across a range of potential solutions, evaluating model performance with four to eight samples based on prior source apportionment literature, PMF identified an optimal solution with 6 factors, or sources. These, corresponding with crustal materials (i.e., soil, dust; Factor 1: V, Si, Ca load highest), secondary ammonium sulfates (Factor 2: SO_4_^2−^, NH_4_^+^, V), biomass burning (Factor 3: Br, K, OC), traffic emissions (Factor 4: Fe, Cu, EC), secondary ammonium nitrates (Factor 5: NH_4_^+^, NO_3_^−^, SO_4_^2−^), and metals potentially from industrial and residential fuel burning (Factor 6: Pb, Zn, Ni, Cu) (Supplementary Figure 2). Distributions of participant source estimates per data acquisition site are displayed in Supplementary Figure 5, while pairwise Spearman correlations between participants’ estimates between sources are provided in Supplementary Figure 6. This approach was performed with exposure estimates from participants across the 21 ABCD Study data collection sites across the U.S., for a data-driven analysis of common, shared PM_2.5_ sources across the nation. These sources are comparable to those found in other published source apportionment studies using data from the U.S. (Rahman and Thurston, 2022; Sarnat et al., 2008).

### 2.5 Analyses

A data analysis plan was registered with the <u>Open Science Framework (OSF)</u> and all code used to run these analyses is available on <u>GitHub</u> (Katie Bottenhorn, 2024). Here, we used our previously published Partial Least Squares Correlation (PLSC; (Sukumaran et al., 2023)) approach to study latent dimensions of associations between exposure to PM_2.5_ sources and cortical microarchitecture.

After excluding individuals based on the previously mentioned MRI quality control criteria (see 2.1), the complete-case sample size for all analyses presented here was 4103 individuals (7334 individuals’ 2-year follow-up visit was prior to March 2020, 6230 individuals’ baseline and 2-year follow-up dMRI data were of sufficient quality). Descriptive statistics were computed across the final dataset (Table 1), including mean and standard deviation for all continuous variables (i.e., age, screen time, physical activity, perceived neighborhood safety, population density, distance to major roadways, nighttime noise, exposure estimates, changes in intracellular diffusion, and head motion); mode, and distribution of all categorical variables (i.e., sex assigned at birth, race and ethnicity, handedness, household income, urbanicity).

**Table 1.** Sample demographics, compared to those of the full ABCD Study.

|  | Full ABCD Study |  | Final Sample* |  |
| --- | --- | --- | --- | --- |
|  | *N* | % | *N* | % |
| Sample size (Total *N*) | 11881 | 100 | 4103 | 34.5% |
| Age at baseline (months) | 118.97 ± 7.50 |  | 119.26 ± 7.45 |  |
| Missing | 41 | <1% | 0 | 0% |
| Time between visits (months) | 24.47 ± 2.32 |  | 23.90 ± 1.68 |  |
| Sex (F) | 5658 | 48% | 1943 | 47% |
| Missing | 41 | <1% | 0 | 0% |
| Handedness (R) | 9398 | 79% | 3287 | 80% |
| Race & Ethnicity |  |  |  |  |
| Asian + Other | 1493 | 13% | 450 | 11% |
| Hispanic | 2405 | 20% | 766 | 19% |
| Black | 1777 | 15% | 405 | 10% |
| White | 6163 | 52% | 2482 | 60% |
| Missing | 43 | <1% | 0 | 0% |
| Household Income |  |  |  |  |
| ≤ \$50,000 | 3215 | 27% | 1735 | 42% |

**Table 1.** Sample demographics, compared to those of the full ABCD Study.
|  | Full ABCD Study |  | Final Sample* |  |
| --- | --- | --- | --- | --- |
|  | *N* | % | *N* | % |
| \$50,000 to \$100,000 | 3066 | 26% | 1921 | 47% |
| ≥ \$100,000 | 4544 | 38% | 2683 | 65% |
| Didn't know or refused | 1013 | 9% | 519 | 13% |
| Missing | 0 | 0% | 0 | 0% |
| Urbanicity |  |  |  |  |
| Urbanized area | 9821 | 83% | 3578 | 87% |
| Urban cluster | 372 | 3% | 157 | 4% |
| Rural area | 966 | 8% | 368 | 9% |
| Missing | 722 | 6% | 0 | 0% |
| MRI Scanner Manufacturer |  |  |  |  |
| Siemens | 7303 | 61% | 2720 | 66% |
| GE Medical Systems | 2941 | 25% | 942 | 23% |
| Philips Medical Systems | 1521 | 13% | 441 | 11% |
| Missing | 116 | 1% | 0 | 0% |
|  | Mean ± Standard deviation |  | Mean ± Standard deviation |  |
| Screen Time (weekday; hours) | 3.46 ± 3.10 |  | 3.11 ± 2.77 |  |
| Screen Time (weekend; hours) | 4.62 ± 3.63 |  | 4.29 ± 3.32 |  |
| Physical Activity (days) | 3.49 ± 2.32 |  | 3.64 ± 2.29 |  |
| Neighborhood Safety | 3.89 ± 0.98 |  | 3.97 ± 0.91 |  |
| Population Density (people/km<sup>2</sup>) | 2136.40 ± 2219.72 |  | 2023.30 ± 2210.08 |  |
| Distance to Roadways (meters) | 1187.58 ± 1282.81 |  | 1218.05 ± 1399.17 |  |
| Nighttime Noise (dB) | 51.10 ± 3.91 |  | 50.69 ± 3.91 |  |
| PM<sub>2.5</sub> (µg/m<sup>3</sup>) | 7.52 ± 2.56 |  | 7.36 ± 2.54 |  |
| Head Motion (mm) | 1.31 ± 0.43 |  | 1.19 ± 0.19 |  |
| Anisotropic diffusion change (%/yr) | -1.28 ± 6.21 |  | -1.00 ± 4.95 |  |
| Isotropic diffusion change (%/yr) | 1.65 ± 3.56 |  | 1.57 ± 3.07 |  |

*Note.* \*Some missing demographic values in the complete-case sample because "complete" was defined as no missing values for variables of interest (i.e., PM<sub>2.5</sub> sources, dMRI data) or covariates (i.e., minimally sufficient set, as described in *Demographic Data, Covariates, & Precision Variables*). The "Other" Race & Ethnicity category includes youth whose caregiver identified them as American Indian/Native American, Alaska Native, Native Hawaiian, Guamanian, Samoan, Other Pacific Islander, Asian Indian, Chinese, Filipino, Japanese, Korean, Vietnamese, Other Asian, Other Race, or as belonging to more than one race.

Prior to performing PSLC, the minimally sufficient set of covariates identified from the DAG were regressed out of both PM_2.5_ source estimates and annualized percent changes in cortical microarchitecture using a linear model implemented in R (v4.2.0), as the PLSC package use here does not accommodate covariate regression. This model included participant age, sex, handedness, race/ethnicity, average physical activity, average weekday and weekend screen time, in addition to data collection site, MRI manufacturer, average head motion during the dMRI scan, household income and their residential address’s proximity to major roadways, urbanicity, area population density, and average nighttime noise level. All following analyses were performed on the residuals from these regressions.

Then, we used PLSC to identify latent dimensions of associations between PM_2.5_ sources and estimates of cortical gray matter intracellular diffusion. To complement and potentially contextualize the PMF-derived PM_2.5_ sources, we also ran PLSC models for each isotropic and anisotropic intracellular diffusion changes with all 15 of the individual chemical components of PM_2.5_. Thus, four PLSC models were run: six PM_2.5_ sources and anisotropic intracellular diffusion, six PM_2.5_ sources and isotropic intracellular diffusion, 15 PM_2.5_ components and anisotropic intracellular diffusion, and 15 PM_2.5_ components and isotropic intracellular diffusion.

Briefly, PLSC is a multivariate, cross-decomposition technique that projects multidimensional variable blocks (here: changes in intracellular diffusion, sources of PM) into a lower dimensional subspace such that the covariance between variable blocks is maximized. This process identifies “latent dimensions” of correspondence between blocks by calculating their best fit correlation, constrained to the maximal covariance structure (Krishnan et al., 2011; McIntosh and Lobaugh, 2004). Because PLSC requires complete case data, we used listwise deletion to remove incomplete cases. Here, we used `TExPosition` to perform PLSC (Beaton et al., 2014), `Boot4PLSC()` from `data4PCCAR` for bootstrapping to identify significant latent dimensions of associations (Abdi and Beaton, 2023). Versions of all packages used in these analyses are noted in Supplementary Methods.

Prior to running PLSC, PM_2.5_ sources or PM_2.5_ components and cortical microarchitecture measures were mean-centered and normalized (to standard deviation of 1). For each block of variables (cortical microarchitecture changes and sources, or components, of PM), the residualized, normalized values were arranged into participant-by-variable matrices. For PM_2.5_ sources, this comprised a row per participant and a column per source, whereas for PM_2.5_ components this included a row per participant and a column per component. For cortical microarchitecture changes, this comprised a row per participant and a column per cortical region. The correlation between the two matrices was then decomposed using singular value decomposition (SVD) into three matrices: a square singular value matrix and a matrix each for PM_2.5_ source (or PM_2.5_ components)and cortical microarchitecture loadings on each latent dimension. Thus, each latent dimension consists of one variable per block of data that was derived from a linear combination of the original variable blocks (i.e., cortical microarchitecture changes, sources or components of PM) and represents the loading of that block on the latent dimension. These variable loadings, or “saliences”, describe how the original variables in each block load contribute to each latent dimension, while the singular values explain the correlation of latent variable pairs, per dimension. An effect size was then calculated per latent dimension, denoting the proportion of covariance between blocks that it explains, from the ratio of that latent dimension’s squared singular values to the sum of all latent dimensions’ singular values.

We, then, used permutation testing and bootstrapping to assess statistical significance of the overall model and of each latent dimension, respectively (Abdi and Williams, 2013; Krishnan et al., 2011; McIntosh and Lobaugh, 2004). In order to test for statistical significance of the overall model, we used the procedure described by McIntosh and Lobaugh (2004), in which variables were reordered across 1000 permutations and the probability of the observed solution was computed as the number of times the permuted singular values exceeded the observed singular values. The SVD used in PLSC gives a fixed number of solutions, the upper bound of which is restricted by the number of variables in the smaller block (in this case, 6 solutions due to 6 sources of air pollution or 15 solutions due to 15 PM_2.5_ components). In order to test for statistical significance of variable loadings on each significant observed latent dimension, we used the procedure described by McIntosh and Lobaugh (2004) and calculated variable loadings from 10,000 bootstrapped samples. These loadings were used to calculate a confidence bootstrap ratio, which approximates a *z*-score, per variable per significant latent dimension. We determined variables with bootstrap ratios greater than 2.5 (i.e., *p* < 0.01) to significantly load on that dimension. This entire process was performed separately for isotropic and anisotropic intracellular diffusion.

Finally, loadings for changes in cortical microarchitecture and sources, or components, of air pollution were plotted against each other to visualize the nature of the relationship. Similarly, the neuroanatomy of each latent dimension of association between cortical microarchitecture development and PM_2.5_ sources or components was visualized via cortical surface plots (i.e., Figure 1C; Supplementary Figures 9C, 10C, 12C, 13C, 14C).

**Figure 1.**
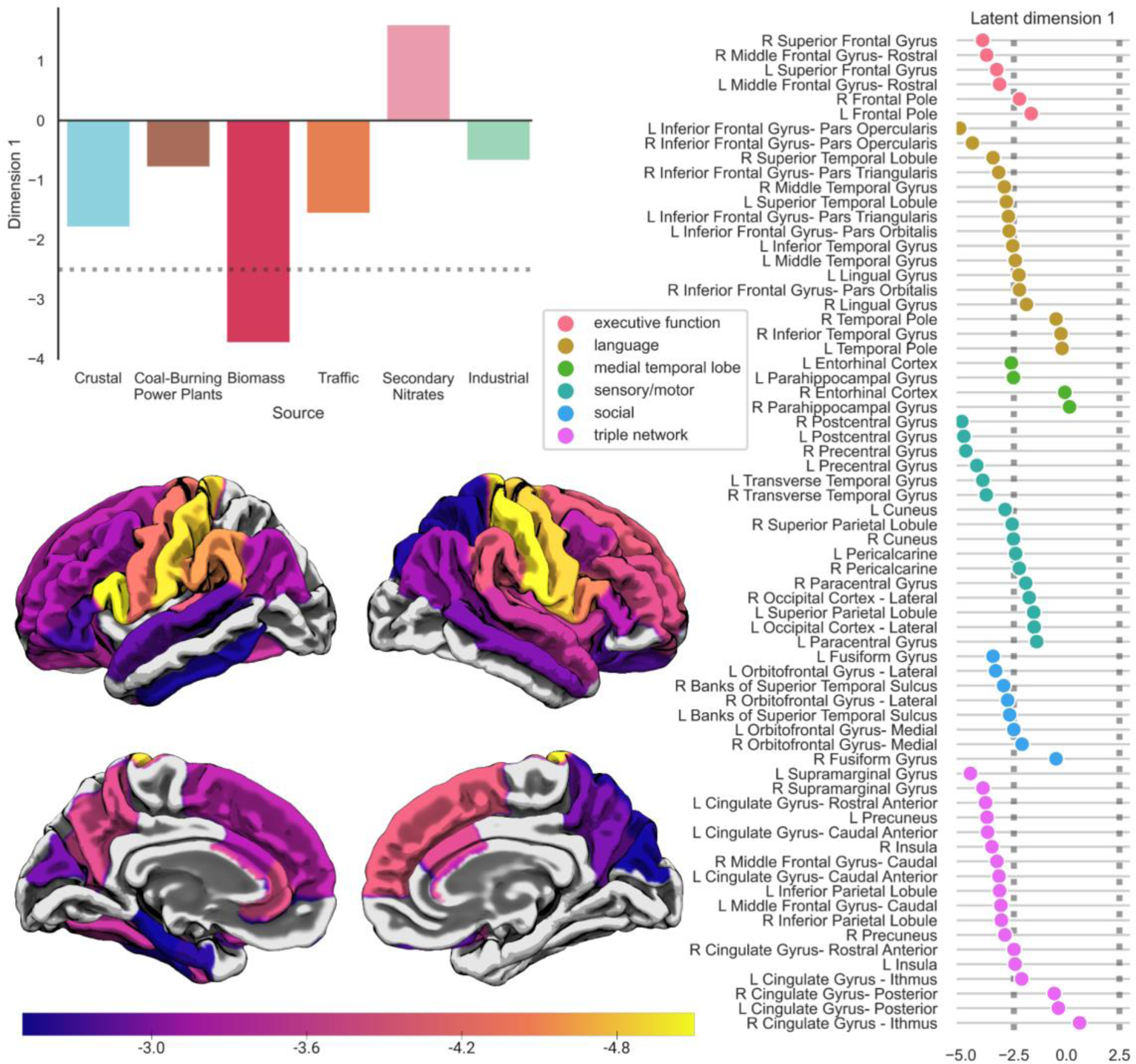
Changes in cortical anisotropic intracellular diffusion are linked to exposure to pollution from biomass burning. A. Bootstrap ratios, equivalent to *z*-scores, indicate significant loadings (|*z*| > 2.5, dashed line) of air pollution from biomass burning on the first latent dimension of brain-pollution associations. B. Bootstrap ratios indicate significant loadings (|*z*| > 2.5; outside the shaded box) of annualized change in anisotropic intracellular diffusion (i.e., neurite density), across brain regions, on the first latent dimension of brain-pollution associations. Regional loading markers are color-coded based on broad functional systems. C. The same regionalloadings (i.e., bootstrap ratios, equivalent to z-scores) of significant changes in anisotropic intracellular diffusion, by region, on the first latent dimension of brain-pollution association, as shown in (B), plotted on a cortical surface template. See Supplementary Table 6 for numerical estimates of brain and pollution saliences.

## 3. Results

The final sample characteristics for the current study are described in Table 1 and average PM_2.5_ component exposures, across participants, are described in Supplementary Table 2 and Supplementary Figure 3. Per-site average participant loadings for each source and average PM_2.5_ component masses are available in Supplementary Table 3 and 4, respectively. Average changes in microarchitecture across the brain in the ABCD Study sample are described elsewhere by Bottenhorn et al. (Bottenhorn et al., 2023).

By assessing latent dimensions of association between PM_2.5_ sources and changes in cortical microarchitecture using PLSC, we revealed a widespread pattern of changes in anisotropic intracellular diffusion related to exposure to PM_2.5_ (*p_omnibus_* < 0.01; Supplementary Table 5), but no significant association between changes in isotropic intracellular diffusion and PM_2.5_ sources (*p_omnibus_* = 0.527; Supplementary Table 5). Analyses of all 15 components of PM_2.5_ and changes in cortical microarchitecture revealed multiple significant latent dimensions of associations between exposure and changes in both anisotropic and isotropic intracellular diffusion (both *p_omniobus_* < 0.01; Supplementary Table 5).

### 3.1 Latent dimensions of association between PM_2.5_ sources, components, and annual changes in anisotropic intracellular diffusion

These analyses revealed one significant latent dimension of association between annual changes in anisotropic intracellular diffusion (i.e., neurite density) and PM_2.5_ sources (*p* < 0.01) that accounted for 68% of shared variance between PM_2.5_ sources and neurite density changes (Supplementary Figure 7, Supplementary Table 5). Exposure to PM_2.5_ attributed to biomass burning (*p* < 0.01; Figure 1A) was associated with neuroanatomically diffuse changes in anisotropic intracellular diffusion (i.e., neurite density) over time, spanning occipital, temporal, parietal, and frontal lobe regions (Figure 1B, C). These exposure-related differences in annual cortical microarchitecture changes were most evident in frontal gyri, pre- and postcentral gyri, superior temporal gyri, inferior parietal regions, caudal anterior cingulate gyri, right insula, and left caudal anterior cingulate gyrus (see Supplementary Table 6 for regional saliences). Recalling that cortical anisotropic intracellular diffusion is predominantly decreasing across this developmental period, these positive exposure-brain associations reflect smaller annual decreases in change in anisotropic intracellular diffusion with age. The remaining latent dimensions were not significant at *p* < 0.01 (Supplementary Table 5).

There were two significant latent dimensions of association between PM_2.5_ components and annualized change in anisotropic intracellular diffusion (both *p* < 0.01) (Supplementary Figure 8). The first dimension represented latent associations between brain development and exposure to elemental carbon (EC), potassium (K), and organic carbon (OC), and ammonium (NH_4_^+^) (Supplementary Figure 9A). These patterns of exposure-related differences reflect smaller annualized change of anisotropic intracellular diffusion and spanned all lobes of the cerebral cortex (Supplementary Figure 9B,C). The second dimension represented latent associations between right hemisphere parahippocampal and fusiform gyri development and exposure to bromine (Br) and copper (Cu), with higher exposure related to greater decreases in annualized changes of anisotropic intracellular diffusion over time (Supplementary Figure 10). The remaining latent dimensions each accounted for <10% of shared variance (see Supplementary Table 5).

### 3.2 Latent dimensions of association between PM_2.5_ sources, components, and annual changes in isotropic intracellular diffusion

While no PM_2.5_ sources showed significant latent dimensions of association with changes to isotropic intracellular diffusion, the PLSC model with PM_2.5_ components identified three latent dimensions of exposure-related differences in isotropic intracellular diffusion change (all p < 0.01; Supplementary Table 5, Supplementary Figure 11). Similar to the PM_2.5_ components and anisotropic intracellular diffusion model, the first dimension represented latent associations between exposure to K, OC, and NH_4_^+^ and brain development (Supplementary Figure 12A). Exposure-related differences in annualized change in isotropic intracellular diffusion along this first dimension was driven by the left cuneus, precentral, middle frontal, and pericalcarine gyri, bilateral anterior cingulate gyri, and right superior frontal gyrus and insula (Supplementary Figure 12B,C). The second dimension represented latent associations between **isotropic intracellular diffusion** brain development and exposure to sulfates (SO_4_^2-^) and vanadium (V) (Supplementary Figure 13A). Exposure-related differences in annual changes in isotropic intracellular diffusion along this second dimension were driven by the right precentral gyrus, bilateral cuneus, and bilateral calcarine gyri (Supplementary Figure 13B,C). Greater exposure to SO_4_ ^2-^ and V was related to greater increases in isotropic intracellular diffusion in cuneus and pericalcarine cortex, but to smaller increases in the prefrontal gyrus. Finally, the third dimension linked vanadium (V) exposure (Supplementary Figure 14A) to differences in in annual isotropic intracellular diffusion changes in superior parietal, lateral occipital, and paracentral gyri, right inferior temporal gyrus, bilateral inferior parietal and right posterior cingulate, insula, and precuneus (Supplementary Figure 14B,C). Again, we see different directions of effect, as greater exposure was associated with greater annual increases in right insula isotropic intracellular diffusion, but less annual increases in parietal, occipital, and inferior temporal isotropic intracellular diffusion changes with age. The remaining, non-significant latent dimensions accounted for <10% each (Supplementary Table 5).

## 4. Discussion

Particulate matter less than 2.5µm in aerodynamic diameter (PM_2.5_) emitted from biomass burning is related to a widespread pattern of cortical microarchitecture development, such that greater exposure is related to smaller changes during early adolescence. Cortical anisotropic intracellular diffusion, thought to reflect neurite (i.e., axon and dendrite) density, is largely decreasing during this developmental period (Bottenhorn et al., 2023), which may reflect ongoing synaptic pruning and subsequent refinement of neural circuitry that occurs during this developmental period. Greater exposure to PM_2.5_ from biomass burning is linked to smaller decreases in neurite density, which may indicate disruption to these ongoing developmental changes. No other source of PM_2.5_ was significantly related to cortical microarchitecture development. However, individual chemical components showed latent dimensions of association with microarchitecture development, suggesting that a single source may have differential effects on brain development due to its specific chemical composition mixture, while individual components might have source-independent effects. Developmental changes in isotropic intracellular diffusion, likely reflecting changing densities of neuronal cell bodies or support cells, was unrelated to source-specific PM exposure. Conversely, individual components of biomass burning (i.e., K, OC) and secondary ammonium sulfates (i.e., NH_4_^+^, SO_4_^2-^, V) were associated both positively and negatively with changes in occipital and frontal regions. This work makes important contributions to the literature on the health effects of PM_2.5_ air pollution exposure in several key ways. First, it assesses exposure-related differences in how the brain *changes* over development, to complement extant cross-sectional neuroimaging studies of brain-exposure associations reviewed in (de Prado Bert et al., 2018; Herting et al., 2019). Second, it assesses *cortical* microarchitecture, using a multi-compartment biophysical model that is sensitive to differences in cortical cytoarchitecture (White et al., 2013), to complement the existing literature on white matter and subcortical microarchitecture (Burnor et al., 2021; Lubczyńska et al., 2020; Peterson et al., 2015; Sukumaran et al., 2023). Third, it leverages a large, geographically diverse sample and PM_2.5_ source apportionment for more interpretable, actionable insights into the role of specific sources of PM_2.5_ exposure in brain development.

### 4.1 Exposures within federal air quality standards linked to differences in cortical microarchitecture changes

The majority of *in vivo* studies of differences in the human brain related to air pollution exposure have been limited by cross-sectional neuroimaging data and are, thus, unable to consider exposure-related brain changes within an individual. Here, we use estimates of annualized change in cortical microarchitecture between ages 9-10 and 11-13 years to capture broad intra-individual change and inter-individual differences therein (Bottenhorn et al., 2023; Mills et al., 2021) during a sensitive period for neurodevelopment (Aoki et al., 2017), including microglial-mediated development (Schalbetter et al., 2022). During this age range, anisotropic intracellular diffusion is decreasing across the cortex (Bottenhorn et al., 2023), but our current results suggest that individuals with greater exposure to PM_2.5_ attributable to biomass burning are decreasing less. Anisotropic, or directional, intracellular diffusion reflects water movement within cylindrical intracellular spaces (e.g., axons, dendrites, glial processes). At the resolution of the diffusion-weighted images used here, anisotropic intracellular diffusion would likely represent aggregated parallel cylindrical intracellular spaces within a voxel (here, 1.7mm isotropic) such as those found in cortical columns and macrocolumns (∼350 to 600µm) (Opris and Casanova, 2014). Thus, the data presented here may provide a coarse estimate of the directional organization of cortical macrocolumns, averaged across a gyrus. In the human brain, synaptic pruning follows a protracted course of development, continuing throughout early adolescence, though decreases in neuronal cell bodies occur earlier in development. The process of synaptic refinement likely includes retraction and fragmentation of neuronal and glial processes (Huttenlocher and Dabholkar, 1997). Anisotropic intracellular diffusion, thought to represent neurite density, may be sensitive to this process. If that is the case, then pruning along an axis (e.g., within a cortical column, in which neurons are vertically connected) could manifest as a decrease in anisotropic intracellular diffusion. This notion is supported by links between anisotropic intracellular diffusion and genes that are responsible for neurite morphogenesis (Brot et al., 2010; Selemon, 2013).

Given the roles of microglia in both healthy developmental synaptic refinement (Schalbetter et al., 2022) and neuroinflammation stemming from particulate exposure (Gillespie et al., 2013; Kang et al., 2021), we should also consider potential roles of microglia in these findings., Interestingly, anisotropic *and* isotropic intracellular diffusion are related to both genes and epigenetic-related cellular processes underlying synaptic pruning and neuroinflammation (Fan et al., 2022). By incorporating *in vivo* neuroimaging with virtual histology and gene expression data from the Allen Brain Atlas, Vidal-Pineiro et al. found that microglia expression was inversely related to cortical thinning in youth, but positively related to cortical thinning in aging adults (Vidal-Pineiro et al., 2020). As cortical thinning is thought to reflect, in part, synaptic pruning, Vidal-Pineiro and colleagues inferred that these contrasting associations reflect a changing role of microglia across the lifespan: from neuronal support in youth to pro-inflammatory immune responses in aging. As our findings might suggest a PM-related difference in the annual decrease in adolescent synaptic pruning seen with age (i.e.,evidenced by a positive association between biomass burning PM and neurite density), is it possible a PM-induced inflammatory response is co-opting the ongoing microglia-mediated synaptic pruning during this period of development? Additional animal research is needed to determine whether the findings presented in this work reflect impaired adolescent cortical pruning as a function of PM exposure.

Experiments in mice suggest that the cortex may have a lower threshold for lasting air pollution-related disruptions than subcortical regions (Bernardi et al., 2021), evidenced by greater increases in microglia and a larger window of antioxidant system susceptibility in cortex, compared to the striatum. These findings highlight the importance of the present work, as the neurotoxicity of air pollution is relatively less investigated in the potentially more susceptible cortex. We found largely positive associations between one-year of exposure and changes in anisotropic intracellular diffusion, suggesting that greater exposure is linked to smaller decreases in neurite density. This is further supported by studies demonstrating air pollution-related differences in cortical macrostructure (i.e., cortical thickness, area, and volume) in similarly-aged children that are consistent with disrupted synaptic pruning (Cserbik et al., 2020; Lubczyńska et al., 2021). More neuroanatomically specific, anisotropic intracellular diffusion is linked to the enrichment of epigenetic markers in the cingulate gyrus (Fan et al., 2022), which is among the regions most impacted by PM_2.5_ exposure seen here and also in rodent models (Nephew et al., 2020). Prior work with the ABCD Study cohort data by our team has revealed that the anterior cingulate gyrus also exhibits largest decreases in anisotropic intracellular diffusion during early adolescence (Bottenhorn et al., 2023). Thus, the relatively strong association between exposure to PM_2.5_ from biomass burning and annual changes to anisotropic intracellular diffusion in the rostral anterior cingulate gyrus may indicate marked regional vulnerability given more dynamic development that occurs in this region during this period of early adolescence.

### 4.2 Source modeling highlights air pollution effects stemming from biomass burning

From nationwide modeled annual estimates of 15 PM_2.5_ components, a positive matrix factorization (PMF) analysis conducted by our research team has identified six overarching sources of PM_2.5_ exposure using exposure profiles from the ABCD Study cohort (Sukumaran et al., Forthcoming). Of these, a source loading highly in bromine (Br), potassium (K), and organic carbon (OC) was attributed to biomass burning (Sukumaran et al., under review). Biomass burning comprises smoke from wildfires, wood burning for residential heating and industrial power production, prescribed burns, land clearing, restaurant emissions (especially from meat cooking), and other sources of organic matter combustion (Adam et al., 2021; Johnston et al., 2019; Li et al., 2018; Robinson et al., 2018; Stockwell et al., 2015). Using these common source factors of PM exposure, our findings add to substantial evidence of specific health effects due to exposure to PM_2.5_ attributed to biomass burning. For example, biomass burning and secondary organic carbon were consistently among the most harmful sources of exposure related to same-day hospital admissions for cardiovascular and respiratory concerns (Krall et al., 2017; Ostro et al., 2007), even when biomass burning only accounted for 7-8% of total PM_2.5_ (Sarnat et al., 2008) cardiovascular, natural, and lung cancer mortality, along with systemic inflammation in Europe (Chen et al., 2022; Huttunen et al., 2012; Siponen et al., 2015) Specifically, the K and OC produced by biomass burning, are consistently associated with overall mortality (Achilleos et al., 2017; Reid et al., 2016). While these previously noted associations are on a different temporal scale and in a broader age range than the data analyzed here, they contextualize our current findings linking individual PM_2.5_ components to brain changes.

Depending on particle size,inhaled PM affects brain health via systemic inflammation following deposition in the lungs, and possibly directly, via the olfactory nerve(Jankowska-Kieltyka et al., 2021; Milton and White, 2020; You et al., 2022). Further, systemic inflammation increases permeability of the blood-brain barrier, potentially allowing circulating ultrafine particles (UFPs; <100nm) in the bloodstream to enter the brain. In vitro human brain models have shown that PM2.5 can cross the blood-brain barrier, accumulate in brain tissue, and cause neuroinflammation and neurodegeneration (Kang et al., 2021). UFPs have also been seen in post mortem brain tissue (Maher et al., 2016), including that of otherwise healthy, young people (Calderón-Garcidueñas et al., 2016, 2010, 2008). Non-human animal studies demonstrate that PM exposure causes oxidative toxicity, lipid peroxidation, cell death, reactive gliosis, and the release of pro-inflammatory cytokines from microglia in the brain (Campbell et al., 2005; Fagundes et al., 2015; Jankowska-Kieltyka et al., 2021; Milton and White, 2020; Scieszka et al., 2022; Woodward et al., 2017; Zhang et al., 2018). Among contributors to PM_2.5_ from biomass burning, wildfire smoke is among the most researched. Wildfire smoke can cause infiltration of peripheral immune cells into the brain, decreases in protective neurometabolites, and increases in pro-inflammatory microglia phenotypes in mice (Scieszka et al., 2022). However, these responses do differ between brain regions (Fagundes et al., 2015; Gerlofs-Nijland et al., 2010; Woodward et al., 2017). Furthermore, prior research has shown that wildfire smoke has a particularly high concentration of UFPs and that PM from wildfire smoke causes greater oxidative stress than PM from other sources (Milton and White, 2020). Altogether, cellular and molecular studies of PM_2.5_ impacts on the brain, especially those of a major biomass burning source in wildfire smoke, underscore the patterns of exposure-related microarchitecture changes uncovered here.

### 4.3 Differences in cortical microarchitecture development coalesce with exposure-related differences in subcortex and white matter at ages 9-10 years

This work fills a gap in the environmental neuroscience literature by assessing changes in cortical microarchitecture related to PM_2.5_ air pollution exposure in humans. Our recent work, also based on the ABCD Study cohort, has identified differences in cortical and subcortical gray matter macrostructure (Cserbik et al., 2020), white matter microarchitecture (Burnor et al., 2021) and subcortical gray matter microarchitecture (Sukumaran et al., 2023) related to total outdoor PM_2.5_ exposure in children ages 9-10 years. Here, we extend this literature with a novel foray assessing differences in *cortical microarchitecture changes related to PM2.5 sources and chemical component exposure*. Notably, we found exposure-related differences in developmental change in the cortical regions connected by the aforementioned white matter tracts affected by PM_2.5_ exposure at ages 9-10 years (e.g., cingulate, frontal, angular, and parietal). Taken together, differences in brain development due to PM_2.5_ exposure may have further implications for adolescent cognitive and mental health. Adolescence is a time of heightened vulnerability to psychopathology, with peak incidence around 14 years of age (Solmi et al., 2022). Abnormal synaptic pruning and exposure to PM_2.5_ have both been linked to increases in mental health problems (Bakolis et al., 2021; Braithwaite et al., 2019; King et al., 2022), along with neurological differences consistent with neurodevelopmental disorders (Xie et al., 2023). Although prior work by our lab and others have found little to no association between total outdoor PM_2.5_ exposure and mental health problems in late childhood and early adolescence (Campbell et al., 2023), these analyses have not been repeated with PM2.5 source or component exposure estimates. A possible explanation, that unifies the neuroimaging and behavioral findings, is that these differences and changes in brain structure and function associated with PM_2.5_ exposure may act as early signs, or biomarkers, of developing psychopathology later on in adolescence and young adulthood. Thus, the differences incortical microarchitecture development related to biomass burning PM_2.5_ exposure shown here may reflect early, pre-clinical impacts with more serious future implications.

### 4.4 Recommendations for parents and policymakers

While PM_2.5_ levels have largely decreased in recent decades (*Our Nation’s Air*, 2022), the contribution from biomass burning has increased in some regions across the US (Milando et al., 2016; Singh et al., 2022)Wildfires are increasingly responsible for PM_2.5_ levels and contribute to biomass burning (Burke et al., 2023; Enayati Ahangar et al., 2021; Li et al., 2021; O’Dell et al., 2019). Further, human-caused climate change and wildfires have a bidirectional relationship, such that climate change is leading to more and larger fires, which release gasses and PM that amplify climate change (Boegelsack et al., 2018). While wildfires are not the only source of biomass burning, wildfire smoke crosses state, province, and national borders, affecting individuals beyond the jurisdictions in which they burn. Thus, exposure to PM_2.5_ from biomass burning is becoming more prevalent even as air quality improves otherwise. Federal and international monitoring and policy-based mitigation efforts are required to lessen the potential impacts of biomass burning on child and adolescent brain development. The demonstrated associations between particles produced by biomass burning and their potential serious threats to human health and child brain development underscore the importance of mitigating exposure. The findings presented here further support the findings of neurological differences in children with exposures below the United States EPA’s national standards, before observable detriment to cognition and mental health. Avoiding exposure, especially in children, will require air quality monitoring, staying indoors when air quality is poor, and frequently changing air filters on air conditioning units or using air filtration to assure good indoor air quality.

### 4.5 Limitations

Geographical location plays a role in the sources and composition of PM_2.5_, but is also linked to many other environmental exposures, both directly and indirectly. Local geography, pollution exposure, built and social environments, sociodemographic factors, and air quality are all intricately linked and can be modeled comprehensively by the “exposome” (Deguen et al., 2022; Pearson et al., 2022; Vineis, 2019). In the current study, we adjusted for many of these sociodemographic and other exposures to try and capture unique associations between one year of PM exposure and brain changes. However, future work should consider a more comprehensive, “omics” approach to partial out potential roles of the built and social environment in brain development, regional variability in exposure profiles, and how they complicate the exposure-neurodevelopment associations described here. Furthermore, PM_2.5_ estimates assessed here represent annual average concentrations from the year 2016 that have been linked to each subject’s primary residence at ages 9-10 years-olds (i.e., the first year of baseline data collection for the ABCD Study). At present, PM_2.5_ levels for subsequent study years are not available for the ABCD Study, precluding an assessment of concurrent exposure and brain changes. However, prior work with these PM_2.5_ data indicates that annual averages are relatively stable in the years leading up to 2016 (Di et al., 2019). More recent data from the U.S. Environmental Protection Agency (EPA) suggests that this stability persisted during the change period assessed here (i.e., from 2016 to March 2020) in most regions across the United States (*Our Nation’s Air*, 2022). Additionally, exposure estimates are linked to participants’ primary addresses and do not include exposure data from other locations at which youth spend time (e.g., school). Thus, while this work highlights potential links between one year of residential exposure and brain changes in the following two years, future work is needed to assess the temporality of these links and incorporate exposures from other frequented locations. Additional years of exposure and neuroimaging data are necessary to clarify how ongoing and cumulative exposures are related to individual differences in neurodevelopmental trajectories.

Here, we used positive matrix factorization (PMF) to perform source apportionment with geographically diverse multi-site PM_2.5_ component estimates (i.e., including data from all 21 ABCD Study data collection, spanning most of the U.S. in one analysis). Doing so enabled us to derive common sources contributing to air pollution levels at all 21 sites. However, this approach also assumes common source profiles across sites, though component profiles may exhibit slight geographical variability within a source. This approach, thus, identifies common sources while potentially overlooking lesser PM_2.5_ sources (e.g., responsible for only a small proportion of PM_2.5_ or only present at a few sites). However, the benefits of deriving common PM_2.5_ outweigh these limitations, as they allow us to study and compare exposure impacts across a large participant sample. The ABCD Study is currently the largest, long-term study of adolescent brain and cognitive development. Thus, it presents a rich opportunity for investigating the health and developmental implications of source-specific air pollution exposure. Prior to running these PLSC models, we adjusted for data collection site as a fixed effect. As some sources identified here exhibit spatial gradients across regions of the U.S., this method of adjusting for site differences may either capture or dilute some sources’ specific effects.

Using region-averaged neuroimaging data simplifies interpretability, reduces dimensionality in a neuroanatomically-informed manner, and may increase signal-to-noise ratio, like the effects of spatial smoothing in MRI data. However, Palmer et al. demonstrate considerable heterogeneity in cortical, subcortical, and white matter microarchitecture within regions using a voxelwise approach (i.e., not averaged across gyri, regions, or tracts) in the same data. On the other hand, the use of PLSC to identify latent dimensions of association between exposure and brain changes limits the interpretability of these findings as it does not provide a readily interpretable effect size with which to contextualize exposure-related differences in change. Future research is needed to assess exposure-neurodevelopment associations seen within regions of cortical microarchitecture and effect sizes of these associations.

Finally, although the ABCD Study sampling procedures aimed to assemble a sample that is sociodemographically representative of the population of the United States, the final sample over represents youth from wealthier, whiter, and more educated households. The quality control procedures employed here and exclusion of individuals with incomplete data further limit generalizability of these findings. Future studies are warranted that include larger representation of lower income and minority backgrounds, which have greater burden to exposure in the U.S. (Collins et al., 2022; Colmer et al., 2020).

## 5. Conclusions

Here, we identified a pattern of differences in early adolescent cortical development linked to exposure to PM_2.5_ air pollution largely attributable to biomass burning. Between ages 9 and 13 years, synaptic pruning is ongoing across the cortex, facilitated in part by the actions of microglia. This process may be reflected by decreasing anisotropic intracellular diffusion, reflecting neurite density, seen across the cortex in this age range. The current study found that individuals with greater PM exposure attributed to biomass burning show smaller decreases in anisotropic intracellular diffusion over time. Policymakers should consider putting greater emphasis on anthropogenic activities that contribute to biomass burning, in order to minimize any long-term harm caused by disrupted cortical development due to PM_2.5_ exposure.

## Acknowledgments

A special thank you to all of the children and families for their participation in their ABCD Study.

Research described in this article was supported by the National Institutes of Health (MMH: NIEHS R01ES032295, R01ES031074; KLB: P30ES07048-27) CCI would like to acknowledge scholars involved in NSP (R25 NS089462), BRAINS (R25 NS094094), and Diversifying CNS (R25 NS117356), as well as R25MH125545 and R25MH120869 for creating a supportive network of ABCD Study users.

Data used in the preparation of this article were obtained from the Adolescent Brain Cognitive Development ^SM^ (ABCD) Study (https://abcdstudy.org), held in the NIMH Data Archive (NDA). This is a multisite, longitudinal study designed to recruit more than 10,000 children age 9-10 and follow them over 10 years into early adulthood. The ABCD Study® is supported by the National Institutes of Health and additional federal partners under award numbers U01DA041048, U01DA050989, U01DA051016, U01DA041022, U01DA051018, U01DA051037, U01DA050987, U01DA041174, U01DA041106, U01DA041117, U01DA041028, U01DA041134, U01DA050988, U01DA051039, U01DA041156, U01DA041025, U01DA041120, U01DA051038, U01DA041148, U01DA041093, U01DA041089, U24DA041123, U24DA041147. A full list of supporters is available at https://abcdstudy.org/federal-partners.html. A listing of participating sites and a complete listing of the study investigators can be found at https://abcdstudy.org/consortium_members/. ABCD consortium investigators designed and implemented the study and/or provided data but did not necessarily participate in the analysis or writing of this report. This manuscript reflects the views of the authors and may not reflect the opinions or views of the NIH or ABCD consortium investigators. The ABCD data repository grows and changes over time. The ABCD data used in this report came from http://dx.doi.org/10.15154/1523041.

## Competing Interests

The authors declare no competing interests.

## Author Contributions

Conceptualization: KLB, MMH, JCC

Data curation: KLB, KS, JS

Formal Analysis: KLB, KS

Funding acquisition: MMH, KLB

Methodology: KLB, MMH, RH, CCI

Project administration: KLB, MMH

Resources: MMH, JS

Software: KLB, KS

Supervision: MMH

Visualization: KLB

Writing – original draft: KLB, MMH

Writing – review & editing: KLB, KS, RH, JS, MMH, JCC

## Notes

### Competing Interest Statement

The authors have declared no competing interest.

### Summary of Updates

Revisions incorporate reviewer comments and include edits to the text and Supplementary Material, but no changes to the analyses or results of this work.

## References

Abdi, H., Beaton, D., 2023. Principal Component and Correspondence Analyses Using R, SpringerBriefs in Statistics. Springer Cham.

Abdi, H., Williams, L.J., 2013. Partial least squares methods: partial least squares correlation and partial least square regression. Comput. Toxicol. Vol. II 549–579.

Achilleos, S., Kioumourtzoglou, M.-A., Wu, C.-D., Schwartz, J.D., Koutrakis, P., Papatheodorou, S.I., 2017. Acute effects of fine particulate matter constituents on mortality: A systematic review and meta-regression analysis. Environ. Int. 109, 89–100. 10.1016/j.envint.2017.09.010

Adam, M.G., Tran, P.T.M., Bolan, N., Balasubramanian, R., 2021. Biomass burning-derived airborne particulate matter in Southeast Asia: A critical review. J. Hazard. Mater. 407, 124760. 10.1016/j.jhazmat.2020.124760

Allen, J.L., Klocke, C., Morris-Schaffer, K., Conrad, K., Sobolewski, M., Cory-Slechta, D.A., 2017a. Cognitive Effects of Air Pollution Exposures and Potential Mechanistic Underpinnings. Curr. Environ. Health Rep. 4, 180–191. 10.1007/s40572-017-0134-3

Allen, J.L., Oberdorster, G., Morris-Schaffer, K., Wong, C., Klocke, C., Sobolewski, M., Conrad, K., Mayer-Proschel, M., Cory-Slechta, D.A., 2017b. Developmental neurotoxicity of inhaled ambient ultrafine particle air pollution: Parallels with neuropathological and behavioral features of autism and other neurodevelopmental disorders. NeuroToxicology 59, 140–154. 10.1016/j.neuro.2015.12.014

Amini, H., Danesh-Yazdi, M., Requia, W., Wei, Y., Abu-Awad, Y., Shi, L., Franklin, M., Kang, C.-M., Wolfson, J., James, P., Habre, R., Zhu, Q., Apte, J., Andersen, Z., Kloog, I., Dominici, F., Koutrakis, P., Schwartz, J., 2022. Hyperlocal super-learned PM2.5 components across the contiguous US. 10.21203/rs.3.rs-1745433/v1

Aoki, C., Romeo, R.D., Smith, S.S., 2017. Adolescence as a critical period for developmental plasticity. Brain Res. 1654, 85–86. 10.1016/j.brainres.2016.11.026

Bakolis, I., Hammoud, R., Stewart, R., Beevers, S., Dajnak, D., MacCrimmon, S., Broadbent, M., Pritchard, M., Shiode, N., Fecht, D., Gulliver, J., Hotopf, M., Hatch, S.L., Mudway, I.S., 2021. Mental health consequences of urban air pollution: prospective population-based longitudinal survey. Soc. Psychiatry Psychiatr. Epidemiol. 56, 1587–1599. 10.1007/s00127-020-01966-x

Barch, D.M., Albaugh, M.D., Avenevoli, S., Chang, L., Clark, D.B., Glantz, M.D., Hudziak, J.J., Jernigan, T.L., Tapert, S.F., Yurgelun-Todd, D., Alia-Klein, N., Potter, A.S., Paulus, M.P., Prouty, D., Zucker, R.A., Sher, K.J., 2018. Demographic, physical and mental health assessments in the adolescent brain and cognitive development study: Rationale and description. Dev. Cogn. Neurosci. 32, 55–66. 10.1016/J.DCN.2017.10.010

Bateson, T.F., Schwartz, J., 2007. Children’s Response to Air Pollutants. J. Toxicol. Environ. Health A 71, 238 –243. 10.1080/15287390701598234

Beaton, D., Chin Fatt, C.R., Abdi, H., 2014. An ExPosition of multivariate analysis with the singular value decomposition in R. Comput. Stat. Data Anal. 72, 176–189. 10.1016/j.csda.2013.11.006

Bekbulat, B., Apte, J.S., Millet, D.B., Robinson, A.L., Wells, K.C., Presto, A.A., Marshall, J.D., 2021. Changes in criteria air pollution levels in the US before, during, and after Covid-19 stay-at-home orders: Evidence from regulatory monitors. Sci. Total Environ. 769, 144693. 10.1016/j.scitotenv.2020.144693

Bernardi, R.B., Zanchi, A.C.T., Damaceno-Rodrigues, N.R., Veras, M.M., Saldiva, P.H.N., Barros, H.M.T., Rhoden, C.R., 2021. The impact of chronic exposure to air pollution over oxidative stress parameters and brain histology. Environ. Sci. Pollut. Res. 28, 47407–47417. 10.1007/s11356-021-14023-0

Binter, A.-C., Kusters, M.S.W., van den Dries, M.A., Alonso, L., Lubczyńska, M.J., Hoek, G., White, T., Iñiguez, C., Tiemeier, H., Guxens, M., 2022. Air pollution, white matter microstructure, and brain volumes: Periods of susceptibility from pregnancy to preadolescence. Environ. Pollut. 313, 120109. 10.1016/j.envpol.2022.120109

Boegelsack, N., Withey, J., O’Sullivan, G., McMartin, D., 2018. A Critical Examination of the Relationship between Wildfires and Climate Change with Consideration of the Human Impact. J. Environ. Prot. 9, 461–467. 10.4236/jep.2018.95028

Bottenhorn, K.L., 2024. 62442katieb/pm2.5sourceXbraindev: Revised code, per Reviews from Environmental International. 10.5281/zenodo.11112232

Bottenhorn, K.L., Cardenas-Iniguez, C., Mills, K.L., Laird, A.R., Herting, M.M., 2023. Profiling intra- and inter-individual differences in brain development across early adolescence. NeuroImage 279, 120287. 10.1016/j.neuroimage.2023.120287

Braithwaite, I., Zhang, S., Kirkbride, J.B., Osborn, D.P.J., Hayes, J.F., 2019. Air Pollution (Particulate Matter) Exposure and Associations with Depression, Anxiety, Bipolar, Psychosis and Suicide Risk: A Systematic Review and Meta-Analysis. Environ. Health Perspect. 127, 126002. 10.1289/EHP4595

Brockmeyer, S., d’Angiulli, A., 2016. How air pollution alters brain development: the role of neuroinflammation. Transl. Neurosci. 7, 24–30.

Brot, S., Rogemond, V., Perrot, V., Chounlamountri, N., Auger, C., Honnorat, J., Moradi-Améli, M., 2010. CRMP5 Interacts with Tubulin to Inhibit Neurite Outgrowth, Thereby Modulating the Function of CRMP2. J. Neurosci. 30, 10639–10654. 10.1523/JNEUROSCI.0059-10.2010

Brumberg, H.L., Karr, C.J., Bole, A., Ahdoot, S., Balk, S.J., Bernstein, A.S., Byron, L.G., Landrigan, P.J., Marcus, S.M., Nerlinger, A.L., Pacheco, S.E., Woolf, A.D., Zajac, L., Baum, C.R., Campbell, C.C., Sample, J.A., Spanier, A.J., Trasande, L., COUNCIL ON ENVIRONMENTAL HEALTH, 2021. Ambient Air Pollution: Health Hazards to Children. Pediatrics 147, e2021051484. 10.1542/peds.2021-051484

Burke, M., Childs, M.L., de la Cuesta, B., Qiu, M., Li, J., Gould, C.F., Heft-Neal, S., Wara, M., 2023. The contribution of wildfire to PM2.5 trends in the USA. Nature 1–6. 10.1038/s41586-023-06522-6

Burnor, E., Cserbik, D., Cotter, D.L., Palmer, C.E., Ahmadi, H., Eckel, S.P., Berhane, K., McConnell, R., Chen, J.-C., Schwartz, J., Jackson, R., Herting, M.M., 2021. Association of Outdoor Ambient Fine Particulate Matter With Intracellular White Matter Microstructural Properties Among Children. JAMA Netw. Open 4, e2138300. 10.1001/jamanetworkopen.2021.38300

Calderón-Garcidueñas, L., Calderón-Garcidueñas, A., Torres-Jardón, R., Avila-Ramírez, J., Kulesza, R.J., Angiulli, A.D., 2015. Air pollution and your brain: what do you need to know right now. Prim. Health Care Res. Dev. 16, 329 –345.

Calderón-Garcidueñas, L., Franco-Lira, M., Henríquez-Roldán, C., Osnaya, N., González-Maciel, A., Reynoso-Robles, R., Villarreal-Calderon, R., Herritt, L., Brooks, D., Keefe, S., 2010. Urban air pollution: influences on olfactory function and pathology in exposed children and young adults. Exp. Toxicol. Pathol. 62, 91–102.

Calderón-Garcidueñas, L., Mora-Tiscareño, A., Styner, M., Gómez-garza, G., Zhu, H., Torres-Jardón, R., Carlos, E., Solorio-López, E., Medina-Cortina, H., Kavanaugh, M., D’Angiulli, A., 2012. White Matter Hyperintensities, Systemic Inflammation, Brain Growth, and Cognitive Functions in Children Exposed to Air Pollution. J. Alzheimers Dis. 31, 183–191. 10.3233/JAD-2012-120610

Calderón-Garcidueñas, L., Reynoso-Robles, R., Vargas-Martínez, J., Gómez-Maqueo-Chew, A., Pérez-Guillé, B., Mukherjee, P.S., Torres-Jardón, R., Perry, G., Gónzalez-Maciel, A., 2016. Prefrontal white matter pathology in air pollution exposed Mexico City young urbanites and their potential impact on neurovascular unit dysfunction and the development of Alzheimer’s disease. Environ. Res. 146, 404–417. 10.1016/j.envres.2015.12.031

Calderón-Garcidueñas, L., Solt, A.C., Henríquez-Roldán, C., Torres-Jardón, R., Nuse, B., Herritt, L., Villarreal-Calderón, R., Osnaya, N., Stone, I., García, R., Brooks, D.M., González-Maciel, A., Reynoso-Robles, R., Delgado-Chávez, R., Reed, W., 2008. Long-term Air Pollution Exposure Is Associated with Neuroinflammation, an Altered Innate Immune Response, Disruption of the Blood-Brain Barrier, Ultrafine Particulate Deposition, and Accumulation of Amyloid β-42 and α-Synuclein in Children and Young Adults. Toxicol. Pathol. 36, 289–310. 10.1177/0192623307313011

Campbell, A., Oldham, M., Becaria, A., Bondy, S.C., Meacher, D., Sioutas, C., Misra, C., Mendez, L.B., Kleinman, M., 2005. Particulate matter in polluted air may increase biomarkers of inflammation in mouse brain. Neurotoxicology 26, 133–140.

Campbell, C.E., Cotter, D.L., Bottenhorn, K.L., Burnor, E., Ahmadi, H., Gauderman, W.J., Cardenas-Iniguez, C., Hackman, D., McConnell, R., Berhane, K., Schwartz, J., Chen, J.-C., Herting, M.M., 2023. Air pollution and age-dependent changes in emotional behavior across early adolescence in the U.S. Environ. Res. 117390. 10.1016/j.envres.2023.117390

Cardenas-Iniguez, C., Burnor, E., Herting, M.M., 2022. Neurotoxicants, the Developing Brain, and Mental Health. Biol. Psychiatry Glob. Open Sci., Exposome: Understanding Environmental Impacts on Brain Development and Risk for Psychopathology 2, 223–232. 10.1016/j.bpsgos.2022.05.002

Carper, R.A., Treiber, J.M., White, N.S., Kohli, J.S., Müller, R.-A., 2017. Restriction Spectrum Imaging As a Potential Measure of Cortical Neurite Density in Autism. Front. Neurosci. 10.

Casey, B.J., Cannonier, T., Conley, M.I., Cohen, A.O., Barch, D.M., Heitzeg, M.M., Soules, M.E., Teslovich, T., Dellarco, D., Garavan, H., Orr, C.A., Wager, T.D., Banich, M.T., Speer, N.K., Sutherland, M.T., Riedel, M.C., Dick, A.S., Bjork, J.M., Thomas, K.M., Chaarani, B., Mejia, M.H., Hagler, D.J., Daniela Cornejo, M., Sicat, C.S., Harms, M.P., Dosenbach, N.U.F., Rosenberg, M., Earl, E., Bartsch, H., Watts, R., Polimeni, J.R., Kuperman, J.M., Fair, D.A., Dale, A.M., 2018. The Adolescent Brain Cognitive Development (ABCD) study: Imaging acquisition across 21 sites. Dev. Cogn. Neurosci. 32, 43–54. 10.1016/J.DCN.2018.03.001

Center for International Earth Science Information Network [WWW Document], n.d. URL http://www.ciesin.org/data.html (accessed 8.25.23).

Chen, C., Warrington, J.A., Dominici, F., Peng, R.D., Esty, D.C., Bobb, J.F., Bell, M.L., 2021. Temporal variation in association between short-term exposure to fine particulate matter and hospitalisations in older adults in the USA: a long-term time-series analysis of the US Medicare dataset. Lancet Planet. Health 5, e534–e541. 10.1016/S2542-5196(21)00168-6

Chen, J., Hoek, G., de Hoogh, K., Rodopoulou, S., Andersen, Z.J., Bellander, T., Brandt, J., Fecht, D., Forastiere, F., Gulliver, J., Hertel, O., Hoffmann, B., Hvidtfeldt, U.A., Verschuren, W.M.M., Jöckel, K.-H., Jørgensen, J.T., Katsouyanni, K., Ketzel, M., Méndez, D.Y., Leander, K., Liu, S., Ljungman, P., Faure, E., Magnusson, P.K.E., Nagel, G., Pershagen, G., Peters, A., Raaschou-Nielsen, O., Rizzuto, D., Samoli, E., van der Schouw, Y.T., Schramm, S., Severi, G., Stafoggia, M., Strak, M., Sørensen, M., Tjønneland, A., Weinmayr, G., Wolf, K., Zitt, E., Brunekreef, B., Thurston, G.D., 2022. Long-Term Exposure to Source-Specific Fine Particles and Mortality—A Pooled Analysis of 14 European Cohorts within the ELAPSE Project. Environ. Sci. Technol. 56, 9277–9290. 10.1021/acs.est.2c01912

Chen, K.L., Henneman, L.R.F., Nethery, R.C., 2021. Differential impacts of COVID-19 lockdowns on PM2.5 across the United States. Environ. Adv. 6, 100122. 10.1016/j.envadv.2021.100122

Chung, Y., Dominici, F., Wang, Y., Coull, B.A., Bell, M.L., 2015. Associations between Long-Term Exposure to Chemical Constituents of Fine Particulate Matter (PM2.5) and Mortality in Medicare Enrollees in the Eastern United States. Environ. Health Perspect. 123, 467–474. 10.1289/ehp.1307549

Clifford, A., Lang, L., Chen, R., Anstey, K.J., Seaton, A., 2016. Exposure to air pollution and cognitive functioning across the life course – A systematic literature review. Environ. Res. 147, 383–398. 10.1016/j.envres.2016.01.018

Collins, T.W., Grineski, S.E., Shaker, Y., Mullen, C.J., 2022. Communities of color are disproportionately exposed to long-term and short-term PM2.5 in metropolitan America. Environ. Res. 214, 114038. 10.1016/j.envres.2022.114038

Colmer, J., Hardman, I., Shimshack, J., Voorheis, J., 2020. Disparities in PM2.5 air pollution in the United States. Science 369, 575–578. 10.1126/science.aaz9353

Compton, W.M., Dowling, G.J., Garavan, H., 2019. Ensuring the Best Use of Data. JAMA Pediatr. 173, 809–810. 10.1001/jamapediatrics.2019.2081

Conley, M.I., Skalaban, L.J., Rapuano, K.M., Gonzalez, R., Laird, A.R., Dick, A.S., Sutherland, M.T., Watts, R., Casey, B.J., 2021. Altered hippocampal microstructure and function in children who experienced Hurricane Irma. Dev. Psychobiol. 63, 864–877. 10.1002/dev.22071

Cory-Slechta, D.A., Merrill, A., Sobolewski, M., 2023. Air Pollution-Related Neurotoxicity Across the Life Span. Annu. Rev. Pharmacol. Toxicol. 63, 143–163. 10.1146/annurev-pharmtox-051921-020812

Costa, L.G., Cole, T.B., Coburn, J., Chang, Y.-C., Dao, K., Roqué, P.J., 2017. Neurotoxicity of traffic-related air pollution. NeuroToxicology 59, 133–139. 10.1016/j.neuro.2015.11.008

Cserbik, D., Chen, J.-C., McConnell, R., Berhane, K., Sowell, E.R., Schwartz, J., Hackman, D.A., Kan, E., Fan, C.C., Herting, M.M., 2020. Fine particulate matter exposure during childhood relates to hemispheric-specific differences in brain structure. Environ. Int. 143, 105933. 10.1016/j.envint.2020.105933

de Prado Bert, P., Mercader, E.M.H., Pujol, J., Sunyer, J., Mortamais, M., 2018. The Effects of Air Pollution on the Brain: a Review of Studies Interfacing Environmental Epidemiology and Neuroimaging. Curr. Environ. Health Rep. 5, 351–364. 10.1007/s40572-018-0209-9

Deguen, S., Amuzu, M., Simoncic, V., Kihal-Talantikite, W., 2022. Exposome and Social Vulnerability: An Overview of the Literature Review. Int. J. Environ. Res. Public. Health 19, 3534. 10.3390/ijerph19063534

Desikan, R.S., Ségonne, F., Fischl, B., Quinn, B.T., Dickerson, B.C., Blacker, D., Buckner, R.L., Dale, A.M., Maguire, R.P., Hyman, B.T., Albert, M.S., Killiany, R.J., 2006. An automated labeling system for subdividing the human cerebral cortex on MRI scans into gyral based regions of interest. NeuroImage 31, 968–980. 10.1016/j.neuroimage.2006.01.021

Di, Q., Amini, H., Shi, L., Kloog, I., Silvern, R., Kelly, J., Sabath, M.B., Choirat, C., Koutrakis, P., Lyapustin, A., Wang, Y., Mickley, L.J., Schwartz, J., 2019. An ensemble-based model of PM2.5 concentration across the contiguous United States with high spatiotemporal resolution. Environ. Int. 130, 104909. 10.1016/j.envint.2019.104909

Echeverria, S.E., Diez-Roux, A.V., Link, B.G., 2004. Reliability of self-reported neighborhood characteristics. J. Urban Health Bull. N. Y. Acad. Med. 81, 682–701. 10.1093/jurban/jth151

Enayati Ahangar, F., Pakbin, P., Hasheminassab, S., Epstein, S.A., Li, X., Polidori, A., Low, J., 2021. Long-term trends of PM2.5 and its carbon content in the South Coast Air Basin: A focus on the impact of wildfires. Atmos. Environ. 255, 118431. 10.1016/j.atmosenv.2021.118431

Essers, E., Binter, A.-C., Neumann, A., White, T., Alemany, S., Guxens, M., 2023. Air pollution exposure during pregnancy and childhood, APOE ε4 status and Alzheimer polygenic risk score, and brain structural morphology in preadolescents. Environ. Res. 216, 114595. 10.1016/j.envres.2022.114595

Fagundes, L.S., Fleck, A. da S., Zanchi, A.C., Saldiva, P.H.N., Rhoden, C.R., 2015. Direct contact with particulate matter increases oxidative stress in different brain structures. Inhal. Toxicol. 27, 462–467.

Fan, C.C., Loughnan, R., Makowski, C., Pecheva, D., Chen, C.-H., Hagler, D.J., Thompson, W.K., Parker, N., van der Meer, D., Frei, O., Andreassen, O.A., Dale, A.M., 2022. Multivariate genome-wide association study on tissue-sensitive diffusion metrics highlights pathways that shape the human brain. Nat. Commun. 13, 2423. 10.1038/s41467-022-30110-3

Fan, C.C., Marshall, A., Smolker, H., Gonzalez, M.R., Tapert, S.F., Barch, D.M., Sowell, E., Dowling, G.J., Cardenas-Iniguez, C., Ross, J., Thompson, W.K., Herting, M.M., 2021. Adolescent Brain Cognitive Development (ABCD) study Linked External Data (LED): Protocol and practices for geocoding and assignment of environmental data. Dev. Cogn. Neurosci. 52, 101030. 10.1016/j.dcn.2021.101030

Ferreira, A.P.S., Ramos, J.M.O., Gamaro, G.D., Gioda, A., Gioda, C.R., Souza, I.C.C., 2022. Experimental rodent models exposed to fine particulate matter (PM2.5) highlighting the injuries in the central nervous system: A systematic review. Atmospheric Pollut. Res. 13, 101407. 10.1016/j.apr.2022.101407

Fonken, L.K., Xu, X., Weil, Z.M., Chen, G., Sun, Q., Rajagopalan, S., Nelson, R.J., 2011. Air pollution impairs cognition, provokes depressive-like behaviors and alters hippocampal cytokine expression and morphology. Mol. Psychiatry 16, 987–995. 10.1038/mp.2011.76

Genc, S., Zadeoglulari, Z., Fuss, S.H., Genc, K., 2012. The Adverse Effects of Air Pollution on the Nervous System. J. Toxicol. 2012, e782462. 10.1155/2012/782462

Gerlofs-Nijland, M.E., van Berlo, D., Cassee, F.R., Schins, R.P., Wang, K., Campbell, A., 2010. Effect of prolonged exposure to diesel engine exhaust on proinflammatory markers in different regions of the rat brain. Part. Fibre Toxicol. 7, 1–10.

Gorham, L.S., Barch, D.M., 2020. White Matter Tract Integrity, Involvement in Sports, and Depressive Symptoms in Children. Child Psychiatry Hum. Dev. 51, 490–501. 10.1007/s10578-020-00960-3

Guxens, M., Garcia-Esteban, R., Giorgis-Allemand, L., Forns, J., Badaloni, C., Ballester, F., Beelen, R., Cesaroni, G., Chatzi, L., de Agostini, M., de Nazelle, A., Eeftens, M., Fernandez, M.F., Fernández-Somoano, A., Forastiere, F., Gehring, U., Ghassabian, A., Heude, B., Jaddoe, V.W.V., Klümper, C., Kogevinas, M., Krämer, U., Larroque, B., Lertxundi, A., Lertxuni, N., Murcia, M., Navel, V., Nieuwenhuijsen, M., Porta, D., Ramos, R., Roumeliotaki, T., Slama, R., Sørensen, M., Stephanou, E.G., Sugiri, D., Tardón, A., Tiemeier, H., Tiesler, C.M.T., Verhulst, F.C., Vrijkotte, T., Wilhelm, M., Brunekreef, B., Pershagen, G., Sunyer, J., 2014. Air Pollution During Pregnancy and Childhood Cognitive and Psychomotor Development: Six European Birth Cohorts. Epidemiology 25, 636–647.

Guxens, M., Lubczyńska, M.J., Muetzel, R.L., Dalmau-Bueno, A., Jaddoe, V.W.V., Hoek, G., van der Lugt, A., Verhulst, F.C., White, T., Brunekreef, B., Tiemeier, H., El Marroun, H., 2018. Air Pollution Exposure During Fetal Life, Brain Morphology, and Cognitive Function in School-Age Children. Biol. Psychiatry 84, 295–303. 10.1016/j.biopsych.2018.01.016

Hagler, D.J., Hatton, S., Cornejo, M.D., Makowski, C., Fair, D.A., Dick, A.S., Sutherland, M.T., Casey, B.J., Barch, D.M., Harms, M.P., Watts, R., Bjork, J.M., Garavan, H.P., Hilmer, L., Pung, C.J., Sicat, C.S., Kuperman, J., Bartsch, H., Xue, F., Heitzeg, M.M., Laird, A.R., Trinh, T.T., Gonzalez, R., Tapert, S.F., Riedel, M.C., Squeglia, L.M., Hyde, L.W., Rosenberg, M.D., Earl, E.A., Howlett, K.D., Baker, F.C., Soules, M., Diaz, J., de Leon, O.R., Thompson, W.K., Neale, M.C., Herting, M., Sowell, E.R., Alvarez, R.P., Hawes, S.W., Sanchez, M., Bodurka, J., Breslin, F.J., Morris, A.S., Paulus, M.P., Simmons, W.K., Polimeni, J.R., van der Kouwe, A., Nencka, A.S., Gray, K.M., Pierpaoli, C., Matochik, J.A., Noronha, A., Aklin, W.M., Conway, K., Glantz, M., Hoffman, E., Little, R., Lopez, M., Pariyadath, V., Weiss, S.R., Wolff-Hughes, D.L., DelCarmen-Wiggins, R., Feldstein Ewing, S.W., Miranda-Dominguez, O., Nagel, B.J., Perrone, A.J., Sturgeon, D.T., Goldstone, A., Pfefferbaum, A., Pohl, K.M., Prouty, D., Uban, K., Bookheimer, S.Y., Dapretto, M., Galvan, A., Bagot, K., Giedd, J., Infante, M.A., Jacobus, J., Patrick, K., Shilling, P.D., Desikan, R., Li, Y., Sugrue, L., Banich, M.T., Friedman, N., Hewitt, J.K., Hopfer, C., Sakai, J., Tanabe, J., Cottler, L.B., Nixon, S.J., Chang, L., Cloak, C., Ernst, T., Reeves, G., Kennedy, D.N., Heeringa, S., Peltier, S., Schulenberg, J., Sripada, C., Zucker, R.A., Iacono, W.G., Luciana, M., Calabro, F.J., Clark, D.B., Lewis, D.A., Luna, B., Schirda, C., Brima, T., Foxe, J.J., Freedman, E.G., Mruzek, D.W., Mason, M.J., Huber, R., McGlade, E., Prescot, A., Renshaw, P.F., Yurgelun-Todd, D.A., Allgaier, N.A., Dumas, J.A., Ivanova, M., Potter, A., Florsheim, P., Larson, C., Lisdahl, K., Charness, M.E., Fuemmeler, B., Hettema, J.M., Maes, H.H., Steinberg, J., Anokhin, A.P., Glaser, P., Heath, A.C., Madden, P.A., Baskin-Sommers, A., Constable, R.T., Grant, S.J., Dowling, G.J., Brown, S.A., Jernigan, T.L., Dale, A.M., 2019. Image processing and analysis methods for the Adolescent Brain Cognitive Development Study. NeuroImage 202, 116091. 10.1016/j.neuroimage.2019.116091

Heeringa, S.G., Berglund, P.A., 2020. A Guide for Population-based Analysis of the Adolescent Brain Cognitive Development (ABCD) Study Baseline Data (preprint). Neuroscience. 10.1101/2020.02.10.942011

Heo, S., Lee, W., Bell, M.L., 2021. Suicide and Associations with Air Pollution and Ambient Temperature: A Systematic Review and Meta-Analysis. Int. J. Environ. Res. Public. Health 18, 7699. 10.3390/ijerph18147699

Herting, M.M., Johnson, C., Mills, K.L., Vijayakumar, N., Dennison, M., Liu, C., Goddings, A.-L., Dahl, R.E., Sowell, E.R., Whittle, S., Allen, N.B., Tamnes, C.K., 2018. Development of subcortical volumes across adolescence in males and females: A multisample study of longitudinal changes. NeuroImage 172, 194–205. 10.1016/j.neuroimage.2018.01.020

Herting, M.M., Kim, R., Uban, K.A., Kan, E., Binley, A., Sowell, E.R., 2017. Longitudinal changes in pubertal maturation and white matter microstructure. Psychoneuroendocrinology 81, 70–79.

Herting, M.M., Sowell, E.R., 2017. Puberty and structural brain development in humans. Front. Neuroendocrinol. 44, 122–137. 10.1016/j.yfrne.2016.12.003

Herting, M.M., Younan, D., Campbell, C.E., Chen, J.-C., 2019. Outdoor Air Pollution and Brain Structure and Function From Across Childhood to Young Adulthood: A Methodological Review of Brain MRI Studies. Front. Public Health 7.

Holguin, F., 2008. Traffic, Outdoor Air Pollution, and Asthma. Immunol. Allergy Clin. North Am., Environmental Factors and Asthma: What We Learned from Epidemiological Studies 28, 577–588. 10.1016/j.iac.2008.03.008

Huttenlocher, P.R., Dabholkar, A.S., 1997. Regional differences in synaptogenesis in human cerebral cortex. J. Comp. Neurol. 387, 167–178. 10.1002/(SICI)1096-9861(19971020)387:2<167::AID-CNE1>3.0.CO;2-Z

Huttunen, K., Siponen, T., Salonen, I., Yli-Tuomi, T., Aurela, M., Dufva, H., Hillamo, R., Linkola, E., Pekkanen, J., Pennanen, A., Peters, A., Salonen, R.O., Schneider, A., Tiittanen, P., Hirvonen, M.-R., Lanki, T., 2012. Low-level exposure to ambient particulate matter is associated with systemic inflammation in ischemic heart disease patients. Environ. Res. 116, 44–51. 10.1016/j.envres.2012.04.004

Huuskonen, M.T., Liu, Q., Lamorie-Foote, K., Shkirkova, K., Connor, M., Patel, A., Montagne, A., Baertsch, H., Sioutas, C., Morgan, T.E., Finch, C.E., Zlokovic, B.V., Mack, W.J., 2021. Air Pollution Particulate Matter Amplifies White Matter Vascular Pathology and Demyelination Caused by Hypoperfusion. Front. Immunol. 12, 785519. 10.3389/fimmu.2021.785519

Jankowska-Kieltyka, M., Roman, A., Nalepa, I., 2021. The Air We Breathe: Air Pollution as a Prevalent Proinflammatory Stimulus Contributing to Neurodegeneration. Front. Cell. Neurosci. 15.

Jin, T., Amini, H., Kosheleva, A., Danesh Yazdi, M., Wei, Y., Castro, E., Di, Q., Shi, L., Schwartz, J., 2022. Associations between long-term exposures to airborne PM2.5 components and mortality in Massachusetts: mixture analysis exploration. Environ. Health 21, 96. 10.1186/s12940-022-00907-2

Johnston, H.J., Mueller, W., Steinle, S., Vardoulakis, S., Tantrakarnapa, K., Loh, M., Cherrie, J.W., 2019. How Harmful Is Particulate Matter Emitted from Biomass Burning? A Thailand Perspective. Curr. Pollut. Rep. 5, 353–377. 10.1007/s40726-019-00125-4

Kang, Y.J., Tan, H.-Y., Lee, C.Y., Cho, H., 2021. An Air Particulate Pollutant Induces Neuroinflammation and Neurodegeneration in Human Brain Models. Adv. Sci. 8, 2101251. 10.1002/advs.202101251

Kazemiparkouhi, F., Honda, T., Eum, K.-D., Wang, B., Manjourides, J., Suh, H.H., 2022. The impact of Long-Term PM2.5 constituents and their sources on specific causes of death in a US Medicare cohort. Environ. Int. 159, 106988. 10.1016/j.envint.2021.106988

King, J.D., Zhang, S., Cohen, A., 2022. Air pollution and mental health: associations, mechanisms and methods. Curr. Opin. Psychiatry 35, 192–199. 10.1097/YCO.0000000000000771

Krishnan, A., Williams, L.J., McIntosh, A.R., Abdi, H., 2011. Partial Least Squares (PLS) methods for neuroimaging: A tutorial and review. NeuroImage, Multivariate Decoding and Brain Reading 56, 455–475. 10.1016/j.neuroimage.2010.07.034

Kusters, M.S.W., Essers, E., Muetzel, R., Ambrós, A., Tiemeier, H., Guxens, M., 2022. Air pollution exposure during pregnancy and childhood, cognitive function, and emotional and behavioral problems in adolescents. Environ. Res. 214, 113891. 10.1016/j.envres.2022.113891

Li, H.Z., Dallmann, T.R., Li, X., Gu, P., Presto, A.A., 2018. Urban Organic Aerosol Exposure: Spatial Variations in Composition and Source Impacts. Environ. Sci. Technol. 52, 415–426. 10.1021/acs.est.7b03674

Li, Y., Tong, D., Ma, S., Zhang, X., Kondragunta, S., Li, F., Saylor, R., 2021. Dominance of Wildfires Impact on Air Quality Exceedances During the 2020 Record-Breaking Wildfire Season in the United States. Geophys. Res. Lett. 48, e2021GL094908. 10.1029/2021GL094908

Liu, F., Liu, C., Liu, Y., Wang, J., Wang, Y., Yan, B., 2023. Neurotoxicity of the air-borne particles: From molecular events to human diseases. J. Hazard. Mater. 457, 131827. 10.1016/j.jhazmat.2023.131827

Liu, Q., Wang, W., Gu, X., Deng, F., Wang, X., Lin, H., Guo, X., Wu, S., 2021. Association between particulate matter air pollution and risk of depression and suicide: a systematic review and meta-analysis. Environ. Sci. Pollut. Res. 28, 9029–9049. 10.1007/s11356-021-12357-3

Lubczyńska, M.J., Muetzel, R.L., El Marroun, H., Basagaña, X., Strak, M., Denault, W., Jaddoe, V.W.V., Hillegers, M., Vernooij, M.W., Hoek, G., White, T., Brunekreef, B., Tiemeier, H., Guxens, M., 2020. Exposure to Air Pollution during Pregnancy and Childhood, and White Matter Microstructure in Preadolescents. Environ. Health Perspect. 128, 027005. 10.1289/EHP4709

Lubczyńska, M.J., Muetzel, R.L., El Marroun, H., Hoek, G., Kooter, I.M., Thomson, E.M., Hillegers, M., Vernooij, M.W., White, T., Tiemeier, H., Guxens, M., 2021. Air pollution exposure during pregnancy and childhood and brain morphology in preadolescents. Environ. Res. 198, 110446. 10.1016/j.envres.2020.110446

Maher, B.A., Ahmed, I.A.M., Karloukovski, V., MacLaren, D.A., Foulds, P.G., Allsop, D., Mann, D.M.A., Torres-Jardón, R., Calderon-Garciduenas, L., 2016. Magnetite pollution nanoparticles in the human brain. Proc. Natl. Acad. Sci. 113, 10797–10801. 10.1073/pnas.1605941113

Martinez-Heras, E., Grussu, F., Prados, F., Solana, E., Llufriu, S., 2021. Diffusion-Weighted Imaging: Recent Advances and Applications. Semin. Ultrasound CT MRI, Advances in Neuroradiology I 42, 490–506. 10.1053/j.sult.2021.07.006

McIntosh, A.R., Lobaugh, N.J., 2004. Partial least squares analysis of neuroimaging data: applications and advances. NeuroImage, Mathematics in Brain Imaging 23, S250–S263. 10.1016/j.neuroimage.2004.07.020

Mennitt, D., Sherrill, K., Fristrup, K., 2014. A geospatial model of ambient sound pressure levels in the contiguous United States. J. Acoust. Soc. Am. 135, 2746–2764. 10.1121/1.4870481

Milando, C., Huang, L., Batterman, S., 2016. Trends in PM2.5 emissions, concentrations and apportionments in Detroit and Chicago. Atmospheric Environ. Oxf. Engl. 1994 129, 197–209. 10.1016/j.atmosenv.2016.01.012

Mills, K.L., Siegmund, K.D., Tamnes, C.K., Ferschmann, L., Wierenga, L.M., Bos, M.G.N., Luna, B., Li, C., Herting, M.M., 2021. Inter-individual variability in structural brain development from late childhood to young adulthood. NeuroImage 242, 118450. 10.1016/j.neuroimage.2021.118450

Milton, L.A., White, A.R., 2020. The potential impact of bushfire smoke on brain health. Neurochem. Int. 139, 104796. 10.1016/j.neuint.2020.104796

Mujahid, M.S., Diez Roux, A.V., Morenoff, J.D., Raghunathan, T., 2007. Assessing the measurement properties of neighborhood scales: from psychometrics to ecometrics. Am. J. Epidemiol. 165, 858–867. 10.1093/aje/kwm040

Nephew, B.C., Nemeth, A., Hudda, N., Beamer, G., Mann, P., Petitto, J., Cali, R., Febo, M., Kulkarni, P., Poirier, G., King, J., Durant, J.L., Brugge, D., 2020. Traffic-related particulate matter affects behavior, inflammation, and neural integrity in a developmental rodent model. Environ. Res. 183, 109242. 10.1016/j.envres.2020.109242

Newbury, J.B., Stewart, R., Fisher, H.L., Beevers, S., Dajnak, D., Broadbent, M., Pritchard, M., Shiode, N., Heslin, M., Hammoud, R., Hotopf, M., Hatch, S.L., Mudway, I.S., Bakolis, I., 2021. Association between air pollution exposure and mental health service use among individuals with first presentations of psychotic and mood disorders: retrospective cohort study. Br. J. Psychiatry 219, 678–685. 10.1192/bjp.2021.119

Newman, B.T., Patrie, J.T., Druzgal, T.J., 2023. An intracellular isotropic diffusion signal is positively associated with pubertal development in white matter. Dev. Cogn. Neurosci. 63, 101301. 10.1016/j.dcn.2023.101301

O’Dell, K., Ford, B., Fischer, E.V., Pierce, J.R., 2019. Contribution of Wildland-Fire Smoke to US PM2.5 and Its Influence on Recent Trends. Environ. Sci. Technol. 53, 1797–1804. 10.1021/acs.est.8b05430

Oldfield, R.C., 1971. The assessment and analysis of handedness: the Edinburgh inventory. Neuropsychologia 9, 97–113.

Opris, I., Casanova, M.F., 2014. Prefrontal cortical minicolumn: from executive control to disrupted cognitive processing. Brain 137, 1863–1875. 10.1093/brain/awt359

Our Nation’s Air, 2022. United States Environmental Protection Agency.

Palmer, C.E., Pecheva, D., Iversen, J.R., Hagler, D.J., Sugrue, L., Nedelec, P., Fan, C.C., Thompson, W.K., Jernigan, T.L., Dale, A.M., 2022. Microstructural development from 9 to 14 years: Evidence from the ABCD Study. Dev. Cogn. Neurosci. 53, 101044. 10.1016/j.dcn.2021.101044

Paulich, K.N., Ross, J.M., Lessem, J.M., Hewitt, J.K., 2021. Screen time and early adolescent mental health, academic, and social outcomes in 9- and 10-year old children: Utilizing the Adolescent Brain Cognitive Development ^SM^ (ABCD) Study. PloS One 16, e0256591. 10.1371/journal.pone.0256591

Paulus, M.P., Squeglia, L.M., Bagot, K., Jacobus, J., Kuplicki, R., Breslin, F.J., Bodurka, J., Morris, A.S., Thompson, W.K., Bartsch, H., Tapert, S.F., 2019. Screen media activity and brain structure in youth: Evidence for diverse structural correlation networks from the ABCD study. NeuroImage 185, 140–153. 10.1016/j.neuroimage.2018.10.040

Pearson, A.L., Shewark, E.A., Burt, S.A., 2022. Associations between neighborhood built, social, or toxicant conditions and child externalizing behaviors in the Detroit metro area: a cross-sectional study of the neighborhood ‘exposome.’ BMC Public Health 22, 1064. 10.1186/s12889-022-13442-z

Peterson, B.S., Bansal, R., Sawardekar, S., Nati, C., Elgabalawy, E.R., Hoepner, L.A., Garcia, W., Hao, X., Margolis, A., Perera, F., Rauh, V., 2022. Prenatal exposure to air pollution is associated with altered brain structure, function, and metabolism in childhood. J. Child Psychol. Psychiatry 63, 1316–1331. 10.1111/jcpp.13578

Peterson, B.S., Rauh, V.A., Bansal, R., Hao, X., Toth, Z., Nati, G., Walsh, K., Miller, R.L., Arias, F., Semanek, D., Perera, F., 2015. Effects of Prenatal Exposure to Air Pollutants (Polycyclic Aromatic Hydrocarbons) on the Development of Brain White Matter, Cognition, and Behavior in Later Childhood. JAMA Psychiatry 72, 531–540. 10.1001/jamapsychiatry.2015.57

Rahman, M.M., Thurston, G., 2022. A hybrid satellite and land use regression model of source-specific PM2.5 and PM2.5 constituents. Environ. Int. 163, 107233. 10.1016/j.envint.2022.107233

Rapuano, K.M., Laurent, J.S., Hagler, D.J., Hatton, S.N., Thompson, W.K., Jernigan, T.L., Dale, A.M., Casey, B.J., Watts, R., 2020. Nucleus accumbens cytoarchitecture predicts weight gain in children. Proc. Natl. Acad. Sci. 117, 26977–26984. 10.1073/pnas.2007918117

Reid, C.E., Brauer, M., Johnston, F.H., Jerrett, M., Balmes, J.R., Elliott, C.T., 2016. Critical Review of Health Impacts of Wildfire Smoke Exposure. Environ. Health Perspect. 124, 1334–1343. 10.1289/ehp.1409277

Robinson, K.M., Price, J.A.B., Demyen, B., 2018. Understanding arithmetic concepts: Does operation matter? J. Exp. Child Psychol. 166, 421–436. 10.1016/j.jecp.2017.09.003

Sarnat, J.A., Marmur, A., Klein, M., Kim, E., Russell, A.G., Sarnat, S.E., Mulholland, J.A., Hopke, P.K., Tolbert, P.E., 2008. Fine Particle Sources and Cardiorespiratory Morbidity: An Application of Chemical Mass Balance and Factor Analytical Source-Apportionment Methods. Environ. Health Perspect. 116, 459–466. 10.1289/ehp.10873

Schalbetter, S.M., von Arx, A.S., Cruz-Ochoa, N., Dawson, K., Ivanov, A., Mueller, F.S., Lin, H.-Y., Amport, R., Mildenberger, W., Mattei, D., Beule, D., Földy, C., Greter, M., Notter, T., Meyer, U., 2022. Adolescence is a sensitive period for prefrontal microglia to act on cognitive development. Sci. Adv. 8, eabi6672. 10.1126/sciadv.abi6672

Scieszka, D., Hunter, R., Begay, J., Bitsui, M., Lin, Y., Galewsky, J., Morishita, M., Klaver, Z., Wagner, J., Harkema, J.R., Herbert, G., Lucas, S., McVeigh, C., Bolt, A., Bleske, B., Canal, C.G., Mostovenko, E., Ottens, A.K., Gu, H., Campen, M.J., Noor, S., 2022. Neuroinflammatory and Neurometabolomic Consequences From Inhaled Wildfire Smoke-Derived Particulate Matter in the Western United States. Toxicol. Sci. Off. J. Soc. Toxicol. 186, 149–162. 10.1093/toxsci/kfab147

Selemon, L.D., 2013. A role for synaptic plasticity in the adolescent development of executive function. Transl. Psychiatry 3, e238–e238. 10.1038/tp.2013.7

Shrier, I., Platt, R.W., 2008. Reducing bias through directed acyclic graphs. BMC Med. Res. Methodol. 8, 70. 10.1186/1471-2288-8-70

Singh, S., Johnson, G., DuBois, D.W., Kavouras, I.G., 2022. Assessment of the Contribution of Local and Regional Biomass Burning on PM2.5 in New York/New Jersey Metropolitan Area. Aerosol Air Qual. Res. 22, 220121. 10.4209/aaqr.220121

Siponen, T., Yli-Tuomi, T., Aurela, M., Dufva, H., Hillamo, R., Hirvonen, M.-R., Huttunen, K., Pekkanen, J., Pennanen, A., Salonen, I., Tiittanen, P., Salonen, R.O., Lanki, T., 2015. Source-specific fine particulate air pollution and systemic inflammation in ischaemic heart disease patients. Occup. Environ. Med. 72, 277–283. 10.1136/oemed-2014-102240

Snider, G., Weagle, C.L., Murdymootoo, K.K., Ring, A., Ritchie, Y., Stone, E., Walsh, A., Akoshile, C., Anh, N.X., Balasubramanian, R., 2016. Variation in global chemical composition of PM 2.5: emerging results from SPARTAN. Atmospheric Chem. Phys. 16, 9629–9653.

Solmi, M., Radua, J., Olivola, M., Croce, E., Soardo, L., Salazar de Pablo, G., Il Shin, J., Kirkbride, J.B., Jones, P., Kim, J.H., Kim, J.Y., Carvalho, A.F., Seeman, M.V., Correll, C.U., Fusar-Poli, P., 2022. Age at onset of mental disorders worldwide: large-scale meta-analysis of 192 epidemiological studies. Mol. Psychiatry 27, 281–295. 10.1038/s41380-021-01161-7

Stockwell, C.E., Veres, P.R., Williams, J., Yokelson, R.J., 2015. Characterization of biomass burning emissions from cooking fires, peat, crop residue, and other fuels with high-resolution proton-transfer-reaction time-of-flight mass spectrometry. Atmospheric Chem. Phys. 15, 845–865. 10.5194/acp-15-845-2015

Sukumaran, K., Bottenhorn, K.L., Schwartz, J., Gauderman, J., Cardenas-Iniguez, C., McConnell, R., Hackman, D.A., Berhane, K., Habre, R., Herting, M.M., Forthcoming. Associations between particulate matter components, their sources, and cognitive outcomes in U.S. children ages 9-10 years-old.

Sukumaran, K., Cardenas-Iniguez, C., Burnor, E., Bottenhorn, K.L., Hackman, D.A., McConnell, R., Berhane, K., Schwartz, J., Chen, J.-C., Herting, M.M., 2023. Ambient fine particulate exposure and subcortical gray matter microarchitecture in 9- and 10-year-old children across the United States. iScience 26, 106087. 10.1016/j.isci.2023.106087

Textor, J., van der Zander, B., Gilthorpe, M.S., Liskiewicz, M., Ellison, G.T., 2016. Robust causal inference using directed acyclic graphs: the R package “dagitty.” Int. J. Epidemiol. 45, 1887–1894. 10.1093/ije/dyw341

Veale, J.F., 2014. Edinburgh handedness inventory–short form: a revised version based on confirmatory factor analysis. Laterality Asymmetries Body Brain Cogn. 19, 164–177.

Vidal-Pineiro, D., Parker, N., Shin, J., French, L., Grydeland, H., Jackowski, A.P., Mowinckel, A.M., Patel, Y., Pausova, Z., Salum, G., Sørensen, Ø., Walhovd, K.B., Paus, T., Fjell, A.M., 2020. Cellular correlates of cortical thinning throughout the lifespan. Sci. Rep. 10, 21803. 10.1038/s41598-020-78471-3

Vineis, P., 2019. What Is the Exposome and How It Can Help Research on Air Pollution. Emiss. Control Sci. Technol. 5, 31–36. 10.1007/s40825-018-0104-8

Volkow, N.D., Koob, G.F., Croyle, R.T., Bianchi, D.W., Gordon, J.A., Koroshetz, W.J., Pérez-Stable, E.J., Riley, W.T., Bloch, M.H., Conway, K., Deeds, B.G., Dowling, G.J., Grant, S., Howlett, K.D., Matochik, J.A., Morgan, G.D., Murray, M.M., Noronha, A., Spong, C.Y., Wargo, E.M., Warren, K.R., Weiss, S.R.B., 2018. The conception of the ABCD study: From substance use to a broad NIH collaboration. Dev. Cogn. Neurosci., The Adolescent Brain Cognitive Development (ABCD) Consortium: Rationale, Aims, and Assessment Strategy 32, 4–7. 10.1016/j.dcn.2017.10.002

Wang, Y., Zhong, Y., Liao, J., Wang, G., 2021. PM2.5-related cell death patterns. Int. J. Med. Sci. 18, 1024–1029. 10.7150/ijms.46421

White, N.S., Leergaard, T.B., D’Arceuil, H., Bjaalie, J.G., Dale, A.M., 2012. Probing tissue microstructure with restriction spectrum imaging: Histological and theoretical validation. Hum. Brain Mapp. 34, 327–346. 10.1002/hbm.21454

White, N.S., McDonald, C., Farid, N., Kuperman, J., Karow, D., Schenker-Ahmed, N.M., Bartsch, H., Rakow-Penner, R., Holland, D., Shabaik, A., Bjørnerud, A., Hope, T., Hattangadi-Gluth, J., Liss, M., Parsons, J.K., Chen, C.C., Raman, S., Margolis, D., Reiter, R.E., Marks, L., Kesari, S., Mundt, A.J., Kane, C.J., Carter, B.S., Bradley, W.G., Dale, A.M., 2014. Diffusion-weighted imaging in cancer: Physical foundations and applications of Restriction Spectrum Imaging. Cancer Res. 74, 4638–4652. 10.1158/0008-5472.CAN-13-3534

White, N.S., McDonald, C.R., Farid, N., Kuperman, J.M., Kesari, S., Dale, A.M., 2013. Improved Conspicuity and Delineation of High-Grade Primary and Metastatic Brain Tumors Using “Restriction Spectrum Imaging”: Quantitative Comparison with High B-Value DWI and ADC. Am. J. Neuroradiol. 34, 958–964. 10.3174/ajnr.A3327

Woodward, N.C., Haghani, A., Johnson, R.G., Hsu, T.M., Saffari, A., Sioutas, C., Kanoski, S.E., Finch, C.E., Morgan, T.E., 2018. Prenatal and early life exposure to air pollution induced hippocampal vascular leakage and impaired neurogenesis in association with behavioral deficits. Transl. Psychiatry 8, 1–10. 10.1038/s41398-018-0317-1

Woodward, N.C., Levine, M.C., Haghani, A., Shirmohammadi, F., Saffari, A., Sioutas, C., Morgan, T.E., Finch, C.E., 2017. Toll-like receptor 4 in glial inflammatory responses to air pollution in vitro and in vivo. J. Neuroinflammation 14, 1–15.

Xie, C., Xiang, S., Shen, C., Peng, X., Kang, J., Li, Y., Cheng, W., He, S., Banaschewski, T., Barker, G.J., Bokde, A.L.W., Bromberg, U., Büchel, C., Desrivières, S., Flor, H., Grigis, A., Garavan, H., Gowland, P., Heinz, A., Ittermann, B., Martinot, J.-L., Martinot, M.-L.P., Nees, F., Orfanos, D.P., Paus, T., Poustka, L., Fröhner, J.H., Smolka, M.N., Walter, H., Whelan, R., Sahakian, B.J., Robbins, T.W., Schumann, G., Jia, T., Feng, J., 2023. A shared neural basis underlying psychiatric comorbidity. Nat. Med. 29, 1232–1242. 10.1038/s41591-023-02317-4

You, R., Ho, Y.-S., Chang, R.C.-C., 2022. The pathogenic effects of particulate matter on neurodegeneration: a review. J. Biomed. Sci. 29, 15. 10.1186/s12929-022-00799-x

Zhang, Q., Li, Q., Ma, J., Zhao, Y., 2018. PM2. 5 impairs neurobehavior by oxidative stress and myelin sheaths injury of brain in the rat. Environ. Pollut. 242, 994–1001.

