## Supplementary Material for "Air pollution from biomass burning disrupts early adolescent cortical microarchitecture development"

This document includes:

- Supplementary Methods
- Supplementary Tables 0 to 6
- Supplementary Figures 1 to 14

#### Supplementary Methods

##### R Packages & Versions

R version 4.2.0 (2022-04-22); platform: aarch64-apple-darwin20 (64-bit); running under: macOS Monterey 12.1

other attached packages:

- RColorBrewer_1.1-3
- psych_2.3.3
- here_1.0.1 readxl_1.4.2
- gridExtra_2.3
- cowplot_1.1.1
- corrplot_0.92
- plyr_1.8.8
- data4PCCAR_0.1.1
- TCA4CATA_0.1.0
- TExPosition_2.6.10.1
- ExPosition_2.8.23
- prettyGraphs_2.1.6
- lubridate_1.9.2
- forcats_1.0.0
- stringr_1.5.0
- dplyr_1.1.2
- purrr_1.0.1
- readr_2.1.4
- tidyr_1.3.0
- tibble_3.2.1
- ggplot2_3.4.2
- tidyverse_2.0.0

loaded via a namespace (and not attached):

- nlme_3.1-157
- bitops_1.0-7
- matrixStats_0.63.0
- fontquiver_0.2.1
- rprojroot_2.0.3
- tools_4.2.0

*Corresponding Author*: **Drs. Katherine Bottenhorn & Megan Herting**, Postdoctoral Scholar & Associate Professor, Department of Population and Public Health Sciences, Keck School of Medicine, University of Southern California, Los Angeles, CA USA |

- utf8_1.2.3
- R6_2.5.1
- KernSmooth_2.23-20
- colorspace_2.1-0
- withr_2.5.0
- mnormt_2.1.1
- tidyselect_1.2.0
- curl_5.0.0
- compiler_4.2.0
- textshaping_0.3.6
- cli_3.6.1
- xml2_1.3.4
- officer_0.6.2
- fontBitstreamVera_0.1.1
- sandwich_3.0-2
- caTools_1.18.2
- scales_1.2.1
- mvtnorm_1.1-3
- askpass_1.1
- systemfonts_1.0.4
- digest_0.6.29
- gfonts_0.2.0
- pkgconfig_2.0.3
- htmltools_0.5.5
- fastmap_1.1.1
- rvg_0.3.2
- rlang_1.1.1
- httpcode_0.3.0
- shiny_1.7.4
- generics_0.1.2
- zoo_1.8-12
- jsonlite_1.8.4
- gtools_3.9.4
- zip_2.3.0
- car_3.0-12
- magrittr_2.0.3
- modeltools_0.2-23
- Matrix_1.4-1
- Rcpp_1.0.8.3
- munsell_0.5.0
- fansi_1.0.4
- abind_1.4-5
- gdtools_0.3.3
- lifecycle_1.0.3
- stringi_1.7.6
- multcomp_1.4-23
- carData_3.0-5
- MASS_7.3-56
- gplots_3.1.3
- parallel_4.2.0
- promises_1.2.0.1
- ggrepel_0.9.3
- crayon_1.5.1
- lattice_0.20-45
- splines_4.2.0
- hms_1.1.3
- pillar_1.9.0
- uuid_1.1-0
- codetools_0.2-18
- stats4_4.2.0
- crul_1.3
- glue_1.6.2
- fontLiberation_0.1.0
- vctrs_0.6.2
- tzdb_0.3.0
- httpuv_1.6.9
- cellranger_1.1.0
- gtable_0.3.3
- openssl_2.0.6
- reshape_0.8.9
- mime_0.12
- coin_1.4-2
- libcoin_1.0-9
- xtable_1.8-4
- later_1.3.1
- ragg_1.2.5
- survival_3.3-1
- timechange_0.2.0
- TH.data_1.1-2
- ellipsis_0.3.2

***Supplementary Table 0.*** Variables by data release

| Variables from ABCD 4.0 Release | Variables from ABCD 5.0 Release |
|---|---|
| - Age<br>- Sex<br>- Handedness<br>- Race/ethnicity<br>- Household income<br>- MRI scanner manufacturer<br>- Data collection site<br>- Screen time<br>- Physical activity | - Nighttime noise<br>- Fine particulate matter ($PM_{2.5}$) chemical components |

- Anisotropic Intracellular Diffusion
- Isotropic Intracellular Diffusion
- Head Motion (mm)
- Average $PM_{2.5}$
- Perceived neighborhood safety
- Population Density
- Urbanicity
- Distance to Roadways

#### Directed Acyclic Graph (DAG)

Using the notion that a confounder must be related to both the exposure (X) and the outcome (Y) (1), we used the software DAGgitty (2) to assist in the creation and interpretation of a DAG for each specific aim of our study. Using color-coded variables and unidirectional arrows we indicate how potential confounding and precision variables are interrelated and their relations to both exposure and outcome, to identify biasing pathways. This graph (Supplementary Figure 1) was completed using field-specific background knowledge, familiarity with published literature, and intuition from authors MMH and JS. Since exposure estimates are based on the child's primary residential address, a potential confounder (c) must predict both the child's residential location (X) and their brain outcome (Y). Using the above causal diagram and the software DAGgitty, we identified and reviewed potential biasing and backdoor (i.e., biasing, noncausal correlations between X and Y) paths among these variables to identify minimal adjustment sets to reduce potential confounding. The minimal adjustment sets identified by our DAG analysis, along with MRI precision variables, are outlined in bold in Supplemental Figure 1 and include: child's age, sex, race/ethnicity, average weekly physical activity, average screen time use, family income, MRI) scanner manufacturer, and their perceived neighborhood quality, population density, urban vs. rural classification of their primary residential location.

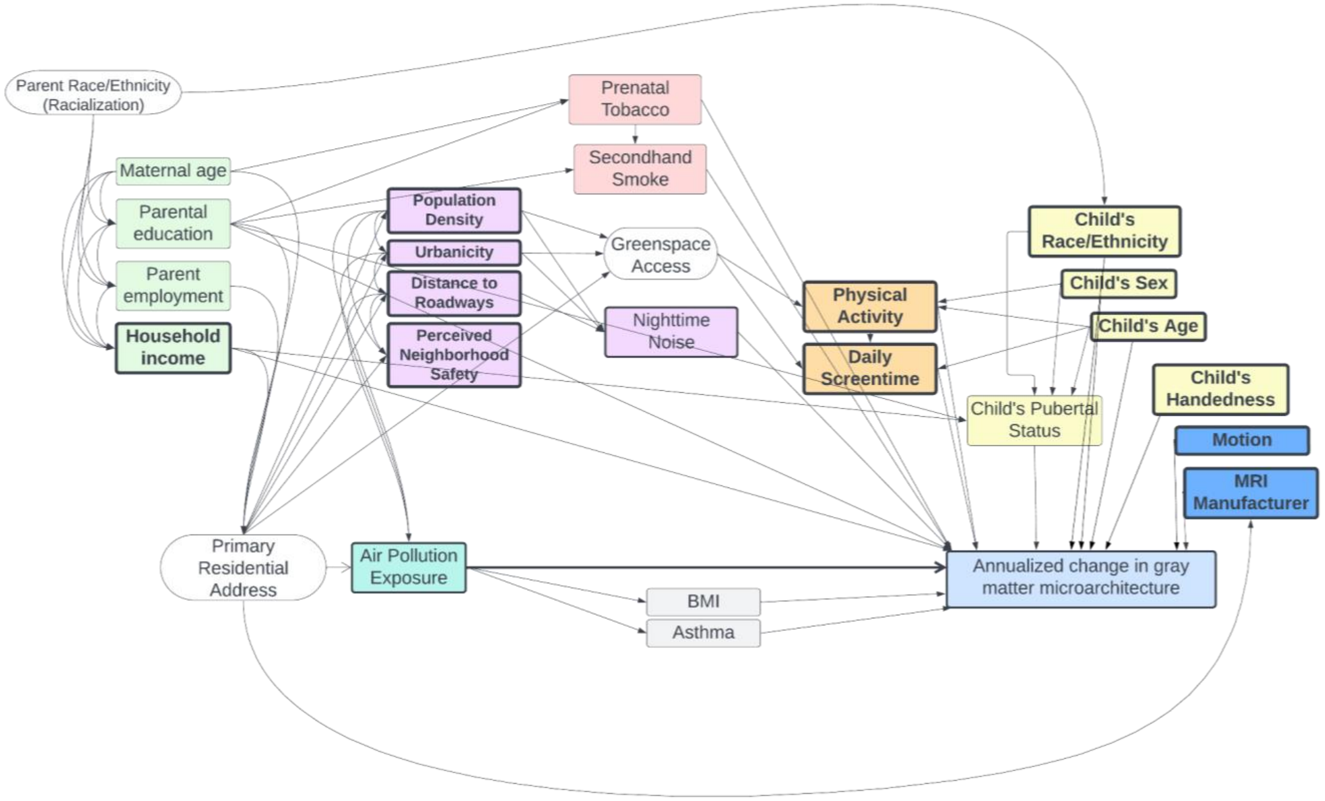


*Supplementary Figure 1*. Directed acyclic graph of factors contributing to air pollution exposure and changes in cortical microstructure. Minimal adjustment set of covariates identified via DAGgitty are bolded. *Abbreviations*: body mass index, BMI; magnetic resonance imaging, MRI.

*Supplementary Table 1. Performance of models predicting PM2.5 component concentrations* (*3*).

| Component | Model RMSE |
|---|---|
| EC | 0.09 |
| $NH_4^+$ | 0.09 |
| $NO_3^-$ | 0.07 |
| OC | 0.18 |
| $SO_4^{2-}$ | 0.28 |
| Br | 0.18 |
| Ca | 2.86 |
| Cu | 1.35 |
| Fe | 4.99 |
| K | 4.77 |
| Ni | 0.45 |
| Pb | 0.34 |
| Si | 10.15 |
| V | 0.73 |
| Zn | 1.82 |

*Abbreviations*: bromine, Br; calcium, Ca; copper, Cu; elemental carbon, EC; iron, Fe; potassium, K; ammonium, $NH_4^+$; nickel, Ni; nitrate, $NO_3^+$; organic carbon, OC; lead, Pb; silicon, Si; sulfate, $SO_4^{2-}$; vanadium, V; zinc, Zn; fine particulate matter <2.5µm, $PM_{2.5}$.

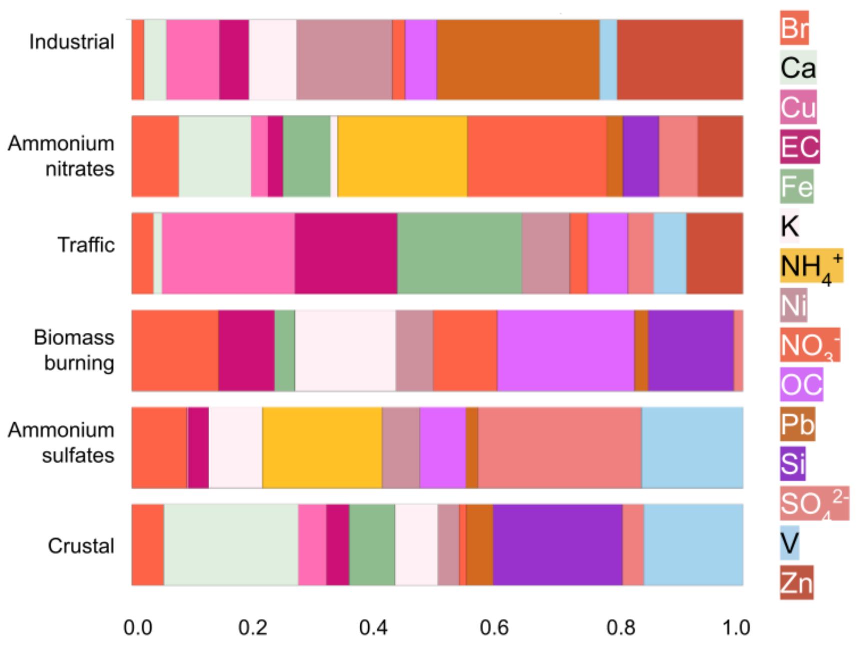


***Supplementary Figure 2.*** Loadings of each $PM_{2.5}$ component on PMF-derived sources of outdoor air pollution. *Abbreviations*: bromine, Br; calcium, Ca; copper, Cu; elemental carbon, EC; iron, Fe; potassium, K; ammonium, $NH_4^+$; nickel, Ni; nitrate, $NO_3^+$; organic carbon, OC; lead, Pb; silicon, Si; sulfate, $SO_4^{2-}$; vanadium, V; zinc, Zn; positive matrix factorization, PMF.

### Supplementary Results

*Supplementary Table 2.* Descriptive statistics of each chemical component of $PM_{2.5}$.

| | Mean | Standard deviation | Minimum | 25th percentile | Median | 75th percentile | Maximum |
|---|---|---|---|---|---|---|---|
| | | | | ng/m³ | | | |
| Br | 2.61 | 0.57 | 1.18 | 2.27 | 2.55 | 2.79 | 5.01 |
| Ca | 48.47 | 21.31 | 11.70 | 30.64 | 45.67 | 63.19 | 118.43 |
| Cu | 4.48 | 1.63 | 0.23 | 3.35 | 4.66 | 5.58 | 8.73 |
| Fe | 63.19 | 23.54 | 12.06 | 47.29 | 61.20 | 77.63 | 166.89 |
| K | 62.58 | 9.58 | 30.68 | 56.09 | 62.92 | 69.45 | 123.68 |
| Ni | 0.81 | 0.26 | 0.01 | 0.71 | 0.84 | 0.93 | 2.14 |
| Pb | 4.49 | 1.26 | 0.60 | 3.84 | 4.70 | 5.32 | 8.22 |
| Si | 84.81 | 38.92 | 24.18 | 52.19 | 71.54 | 115.33 | 231.11 |
| V | 0.37 | 0.20 | 0.01 | 0.25 | 0.34 | 0.46 | 1.26 |
| Zn | 8.96 | 3.78 | 1.25 | 6.79 | 8.57 | 10.67 | 27.17 |
| | | | | μg/m³ | | | |
| EC | 0.51 | 0.15 | 0.11 | 0.41 | 0.50 | 0.59 | 1.41 |
| $NH_4^+$ | 0.28 | 0.12 | 0.05 | 0.20 | 0.28 | 0.37 | 0.80 |
| $NO_3^-$ | 0.92 | 0.34 | 0.14 | 0.62 | 0.93 | 1.17 | 2.64 |
| OC | 1.84 | 0.44 | 0.75 | 1.54 | 1.81 | 2.07 | 3.52 |
| $SO_4^{2-}$ | 0.88 | 0.28 | 0.33 | 0.58 | 0.94 | 1.09 | 2.00 |

*Abbreviations*: bromine, Br; calcium, Ca; copper, Cu; elemental carbon, EC; iron, Fe; potassium, K; ammonium, $NH_4^+$; nickel, Ni; nitrate, $NO_3^+$; organic carbon, OC; lead, Pb; silicon, Si; sulfate, $SO_4^{2-}$; vanadium, V; zinc, Zn; fine particulate matter <2.5μm, $PM_{2.5}$.

*mentary Table 3*. Average and standard deviation of participant contributions to each $PM_{2.5}$ source by data collection site.

| Region | Site | Crustal | Ammonium Sulfates | Biomass Burning | Traffic | Ammonium Nitrates | Industrial |
|---|---|---|---|---|---|---|---|
| West | CHLA | 0.99 ± 0.43 | 1.15 ± 0.29 | 1.76 ± 0.2 | 1.89 ± 0.39 | 1.97 ± 0.53 | 0.61 ± 0.23 |
| | CUB | 1.78 ± 0.33 | 0.14 ± 0.19 | 1.12 ± 0.15 | 0.96 ± 0.37 | 0.87 ± 0.26 | 0.65 ± 0.3 |
| | UTAH | 1.88 ± 0.4 | 0.02 ± 0.12 | 1.28 ± 0.16 | 0.88 ± 0.27 | 1.47 ± 0.2 | 0.99 ± 0.2 |
| | SRI | 0.51 ± 0.16 | 0.61 ± 0.13 | 1.47 ± 0.16 | 1.3 ± 0.2 | 0.81 ± 0.2 | 1.0 ± 0.14 |
| | UCLA | 1.13 ± 0.49 | 1.16 ± 0.29 | 1.71 ± 0.18 | 1.76 ± 0.31 | 1.58 ± 0.46 | 0.69 ± 0.2 |
| | UCSD | 1.21 ± 0.16 | 0.96 ± 0.18 | 1.38 ± 0.18 | 1.41 ± 0.32 | 1.05 ± 0.18 | 0.91 ± 0.22 |
| | OHSU | 0.32 ± 0.11 | 0.42 ± 0.05 | 1.48 ± 0.21 | 1.18 ± 0.34 | 0.25 ± 0.08 | 1.21 ± 0.32 |
| Southwest | LIBR | 2.01 ± 0.26 | 1.36 ± 0.15 | 0.88 ± 0.08 | 0.7 ± 0.25 | 0.78 ± 0.19 | 0.88 ± 0.14 |
| Midwest | UMICH | 0.18 ± 0.19 | 1.11 ± 0.14 | 0.77 ± 0.13 | 0.8 ± 0.42 | 1.75 ± 0.17 | 1.27 ± 0.19 |
| | UMN | 0.41 ± 0.28 | 0.76 ± 0.12 | 0.82 ± 0.11 | 0.54 ± 0.34 | 1.47 ± 0.17 | 1.16 ± 0.31 |
| | UWM | 0.33 ± 0.19 | 0.91 ± 0.14 | 0.7 ± 0.07 | 0.92 ± 0.23 | 1.78 ± 0.16 | 1.51 ± 0.16 |
| | WUSTL | 1.11 ± 0.22 | 1.33 ± 0.15 | 0.75 ± 0.16 | 0.75 ± 0.41 | 1.24 ± 0.19 | 1.15 ± 0.29 |
| Northeast | ROC | 0.13 ± 0.12 | 1.1 ± 0.11 | 0.75 ± 0.11 | 0.53 ± 0.16 | 1.23 ± 0.14 | 1.11 ± 0.12 |
| | UMB | 0.42 ± 0.14 | 1.41 ± 0.15 | 0.78 ± 0.11 | 1.17 ± 0.36 | 0.94 ± 0.14 | 1.23 ± 0.19 |
| | UPMC | 0.47 ± 0.1 | 1.84 ± 0.13 | 0.38 ± 0.09 | 1.67 ± 0.31 | 1.19 ± 0.13 | 1.95 ± 0.17 |
| | UVM | 0.33 ± 0.14 | 0.79 ± 0.1 | 0.91 ± 0.11 | 0.23 ± 0.2 | 0.51 ± 0.12 | 0.67 ± 0.26 |
| | YALE | 0.43 ± 0.15 | 1.12 ± 0.2 | 0.79 ± 0.13 | 0.95 ± 0.47 | 0.64 ± 0.17 | 1.32 ± 0.15 |
| | MUSC | 1.16 ± 0.13 | 1.61 ± 0.16 | 1.17 ± 0.1 | 0.71 ± 0.21 | -0.04 ± 0.1 | 0.73 ± 0.28 |

| | | | | | | | |
|---|---|---|---|---|---|---|---|
| Southeast | FIU | 2.31 ± 0.42 | 1.33 ± 0.17 | 0.43 ± 0.16 | 1.17 ± 0.21 | 0.38 ± 0.17 | 0.47 ± 0.17 |
| | UFL | 1.8 ± 0.19 | 1.48 ± 0.12 | 1.22 ± 0.13 | 0.63 ± 0.23 | -0.06 ± 0.1 | 0.25 ± 0.24 |
| | VCU | 0.45 ± 0.11 | 1.52 ± 0.17 | 0.98 ± 0.16 | 0.79 ± 0.37 | 0.59 ± 0.21 | 0.91 ± 0.36 |

*Abbreviations*: Children's Hospital Los Angeles (CHLA), University of Colorado Boulder (CUB), University of Utah (UTAH), SRI International (SRI), University of California Los Angeles (UCLA), University of California San Diego (UCSD), Oregon Health & Science University (OHSU), Laureate Institute for Brain Research (LIBR), University of Michigan (UMICH), University of Minnesota (UMN), University of Wisconsin-Milwaukee (UWM), Washington University in St. Louis (WUSTL), University of Rochester (ROC), University of Maryland at Baltimore (UMB), University of Pittsburgh (UPMC), University of Vermont (UVM), Yale University (YALE), Medical University of South Carolina (MUSC), Florida International University (FIU), University of Florida (UFL), Virginia Commonwealth University (VCU), fine particulate matter <2.5μm ($PM_{2.5}$).

***Supplementary Table 4***. Average and standard deviation of each $PM_{2.5}$ component mass by data collection site.

| Region | Site | Br | Ca | Cu | EC | Fe | K | $NH_4^+$ | Ni | $NO_3^-$ | OC | Pb | Si | $SO_4^{2-}$ | V | Zn |
|---|---|---|---|---|---|---|---|---|---|---|---|---|---|---|---|---|
| W | CHLA | 3.94 ± 0.41 | 62.87 ± 9.85 | 6.53 ± 0.71 | 0.85 ± 0.19 | 114.44 ± 18.71 | 77.69 ± 9.86 | 0.5 ± 0.11 | 0.88 ± 0.22 | 1.62 ± 0.31 | 2.93 ± 0.29 | 3.74 ± 0.74 | 118.47 ± 25.02 | 1.28 ± 0.21 | 0.28 ± 0.16 | 12.53 ± 5.01 |
| | CUB | 2.03 ± 0.23 | 57.65 ± 9.94 | 4.07 ± 1.23 | 0.53 ± 0.14 | 76.52 ± 16.51 | 58.15 ± 8.64 | 0.19 ± 0.04 | 0.69 ± 0.21 | 0.94 ± 0.19 | 1.62 ± 0.25 | 3.73 ± 1.17 | 155.86 ± 27.78 | 0.53 ± 0.08 | 0.28 ± 0.08 | 6.59 ± 1.91 |
| | UTAH | 3.23 ± 0.52 | 78.42 ± 14.64 | 4.64 ± 0.83 | 0.56 ± 0.1 | 78.94 ± 13.26 | 63.74 ± 6.81 | 0.24 ± 0.04 | 0.8 ± 0.12 | 1.21 ± 0.12 | 1.77 ± 0.2 | 4.99 ± 0.77 | 136.51 ± 12.99 | 0.5 ± 0.04 | 0.36 ± 0.07 | 10.13 ± 1.86 |
| | SRI | 2.59 ± 0.23 | 34.94 ± 3.82 | 5.43 ± 0.55 | 0.56 ± 0.08 | 68.32 ± 9.7 | 64.79 ± 6.77 | 0.18 ± 0.06 | 1.05 ± 0.18 | 0.98 ± 0.14 | 2.29 ± 0.22 | 4.13 ± 0.41 | 60.18 ± 10.23 | 0.66 ± 0.07 | 0.37 ± 0.17 | 8.78 ± 1.21 |
| | UCLA | 3.82 ± 0.39 | 60.7 ± 11.26 | 6.49 ± 0.65 | 0.77 ± 0.14 | 108.14 ± 17.35 | 76.38 ± 6.56 | 0.44 ± 0.09 | 0.95 ± 0.23 | 1.41 ± 0.28 | 2.88 ± 0.26 | 4.1 ± 0.58 | 114.29 ± 23.76 | 1.19 ± 0.18 | 0.34 ± 0.19 | 10.03 ± 3.61 |
| | UCSD | 3.14 ± 0.37 | 54.29 ± 6.5 | 6.01 ± 0.86 | 0.63 ± 0.12 | 83.93 ± 14.57 | 72.08 ± 6.61 | 0.29 ± 0.05 | 1.01 ± 0.21 | 1.1 ± 0.14 | 2.32 ± 0.23 | 4.44 ± 0.78 | 100.81 ± 6.9 | 0.98 ± 0.09 | 0.44 ± 0.12 | 9.03 ± 1.7 |
| | OHSU | 2.1 ± 0.19 | 21.46 ± 4.3 | 5.04 ± 1.17 | 0.55 ± 0.09 | 53.44 ± 16.03 | 66.22 ± 7.87 | 0.09 ± 0.02 | 0.68 ± 0.29 | 0.61 ± 0.04 | 2.25 ± 0.24 | 4.56 ± 1.03 | 48.64 ± 5.52 | 0.46 ± 0.02 | 0.31 ± 0.14 | 10.06 ± 2.94 |
| SW | LIBR | 2.56 ± 0.15 | 76.63 ± 11.94 | 4.12 ± 0.99 | 0.47 ± 0.08 | 63.18 ± 11.62 | 72.21 ± 3.97 | 0.33 ± 0.04 | 0.78 ± 0.11 | 0.77 ± 0.13 | 1.94 ± 0.15 | 4.53 ± 0.53 | 118.83 ± 10.92 | 1.15 ± 0.07 | 0.43 ± 0.07 | 7.94 ± 1.34 |
| MW | UMICH | 2.55 ± 0.2 | 36.0 ± 7.66 | 4.35 ± 1.52 | 0.46 ± 0.09 | 57.84 ± 16.44 | 57.36 ± 4.47 | 0.45 ± 0.05 | 0.86 ± 0.16 | 1.3 ± 0.09 | 1.59 ± 0.17 | 5.24 ± 0.58 | 52.48 ± 4.53 | 0.99 ± 0.08 | 0.31 ± 0.1 | 10.98 ± 2.92 |
| | UMN | 2.39 ± | 36.79 ± 9.11 | 3.93 ± | 0.37 ± 0.07 | 46.42 ± 13.59 | 54.71 ± 4.62 | 0.32 ± 0.04 | 0.74 ± 0.22 | 1.16 ± 0.11 | 1.39 ± 0.16 | 4.88 ± 1.07 | 55.74 ± | 0.75 ± 0.06 | 0.28 ± 0.08 | 8.94 ± 2.09 |

| | | | | | | | | | | | | | | | | |
|---|---|---|---|---|---|---|---|---|---|---|---|---|---|---|---|---|
| | | 0.26 | | 1.43 | | | | | | | | | 10.28 | | | |
| | UWM | 2.56 ± 0.2 | 43.23 ± 6.64 | 5.26 ± 0.97 | 0.44 ± 0.05 | 66.5 ± 9.94 | 58.31 ± 4.09 | 0.43 ± 0.04 | 1.0 ± 0.1 | 1.3 ± 0.1 | 1.63 ± 0.1 | 5.91 ± 0.47 | 52.87 ± 4.87 | 0.89 ± 0.1 | 0.29 ± 0.06 | 12.78 ± 1.92 |
| | WUSTL | 2.6 ± 0.16 | 62.58 ± 8.52 | 4.21 ± 1.55 | 0.47 ± 0.09 | 58.37 ± 15.72 | 63.32 ± 5.06 | 0.4 ± 0.06 | 0.78 ± 0.19 | 0.96 ± 0.1 | 1.84 ± 0.18 | 5.06 ± 0.95 | 71.13 ± 8.35 | 1.1 ± 0.08 | 0.55 ± 0.15 | 10.39 ± 2.8 |
| NE | ROC | 2.44 ± 0.19 | 26.09 ± 3.43 | 3.09 ± 0.69 | 0.36 ± 0.03 | 40.95 ± 8.01 | 47.1 ± 3.83 | 0.3 ± 0.04 | 0.78 ± 0.1 | 0.97 ± 0.11 | 1.47 ± 0.1 | 4.69 ± 0.46 | 43.83 ± 5.08 | 0.91 ± 0.05 | 0.2 ± 0.05 | 8.37 ± 1.16 |
| | UMB | 2.41 ± 0.19 | 33.52 ± 6.56 | 4.93 ± 1.28 | 0.56 ± 0.09 | 62.04 ± 14.54 | 56.72 ± 5.39 | 0.36 ± 0.04 | 0.92 ± 0.1 | 0.86 ± 0.09 | 1.96 ± 0.18 | 5.25 ± 0.57 | 51.62 ± 5.0 | 1.1 ± 0.07 | 0.28 ± 0.09 | 9.76 ± 2.97 |
| | UPMC | 2.83 ± 0.2 | 39.9 ± 4.02 | 6.46 ± 1.03 | 0.68 ± 0.07 | 84.8 ± 12.85 | 64.12 ± 3.17 | 0.44 ± 0.04 | 1.16 ± 0.09 | 0.9 ± 0.08 | 1.97 ± 0.1 | 7.17 ± 0.43 | 57.68 ± 4.19 | 1.45 ± 0.08 | 0.35 ± 0.06 | 21.15 ± 3.74 |
| | UVM | 1.65 ± 0.1 | 20.75 ± 4.04 | 1.72 ± 0.63 | 0.27 ± 0.08 | 21.9 ± 7.5 | 47.92 ± 5.37 | 0.19 ± 0.04 | 0.37 ± 0.3 | 0.54 ± 0.07 | 1.35 ± 0.16 | 2.81 ± 0.9 | 35.94 ± 10.71 | 0.59 ± 0.05 | 0.08 ± 0.05 | 5.32 ± 1.33 |
| | YALE | 2.2 ± 0.16 | 28.22 ± 6.61 | 3.94 ± 1.26 | 0.52 ± 0.14 | 49.93 ± 16.49 | 54.2 ± 4.43 | 0.23 ± 0.04 | 1.02 ± 0.31 | 0.7 ± 0.13 | 1.71 ± 0.24 | 5.23 ± 0.43 | 50.25 ± 6.57 | 0.89 ± 0.08 | 0.28 ± 0.14 | 9.75 ± 2.42 |
| SE | MUSC | 2.65 ± 0.16 | 32.94 ± 4.48 | 3.06 ± 0.71 | 0.42 ± 0.08 | 47.78 ± 7.21 | 65.21 ± 3.69 | 0.2 ± 0.04 | 0.93 ± 0.23 | 0.31 ± 0.05 | 2.06 ± 0.13 | 3.6 ± 0.99 | 87.87 ± 6.74 | 1.07 ± 0.09 | 0.69 ± 0.16 | 5.3 ± 1.56 |
| | FIU | 2.64 ± 0.15 | 66.11 ± 10.33 | 6.22 ± 1.02 | 0.52 ± 0.08 | 71.94 ± 8.24 | 69.48 ± 2.99 | 0.15 ± 0.02 | 0.75 ± 0.15 | 0.54 ± 0.12 | 1.1 ± 0.1 | 3.23 ± 0.75 | 102.36 ± 20.84 | 1.06 ± 0.11 | 0.84 ± 0.16 | 4.21 ± 0.71 |
| | UFL | 2.68 ± 0.17 | 46.14 ± 4.97 | 2.95 ± 1.12 | 0.4 ± 0.08 | 49.42 ± 7.01 | 70.53 ± 3.42 | 0.17 ± 0.05 | 0.64 ± 0.22 | 0.3 ± 0.04 | 1.85 ± 0.16 | 2.14 ± 0.89 | 119.13 ± 11.06 | 1.04 ± 0.06 | 0.5 ± 0.13 | 3.62 ± 0.47 |
| | VCU | 2.43 ± 0.14 | 26.37 ± 3.84 | 3.7 ± 1.84 | 0.44 ± 0.09 | 45.68 ± 11.77 | 56.49 ± 5.14 | 0.32 ± 0.06 | 0.7 ± 0.27 | 0.64 ± 0.1 | 2.01 ± 0.23 | 4.21 ± 1.35 | 55.3 ± 5.23 | 1.08 ± 0.1 | 0.26 ± 0.09 | 6.8 ± 1.74 |

*Note*. Values indicate mean ± standard deviation. Measurements of EC, $NH_4^+$, $NO_3^-$, OC, and $SO_4^{2-}$ are in μg/m$^3$, while measurements of Br, Ca, Cu, Fe, K, Ni, Pb, Si, V, and Zn are in ng/m$^3$. *Abbreviations*: West, W; Southwest, SW; Midwest, MW; Northeast, NE; Southeast, SE; bromine, Br; calcium, Ca; copper, Cu; elemental carbon, EC; iron, Fe; potassium, K; ammonium, $NH_4^+$; nickel, Ni; nitrate, $NO_3^+$; organic carbon, OC; lead, Pb; silicon, Si; sulfate, $SO_4^{2-}$; vanadium, V; zinc, Zn. Children's Hospital Los Angeles (CHLA), University of Colorado Boulder (CUB), University of Utah (UTAH), SRI International (SRI), University of California Los Angeles (UCLA), University of California San Diego (UCSD), Oregon Health & Science University (OHSU), Laureate Institute for Brain Research (LIBR), University of Michigan (UMICH), University of Minnesota (UMN), University of Wisconsin-Milwaukee (UWM), Washington University in St. Louis (WUSTL), University of Rochester (ROC), University of Maryland at Baltimore (UMB), University of Pittsburgh (UPMC), University of Vermont (UVM), Yale University (YALE), Medical University of South Carolina (MUSC), Florida International University (FIU), University of Florida (UFL), Virginia Commonwealth University (VCU), fine particulate matter <2.5μm ($PM_{2.5}$).

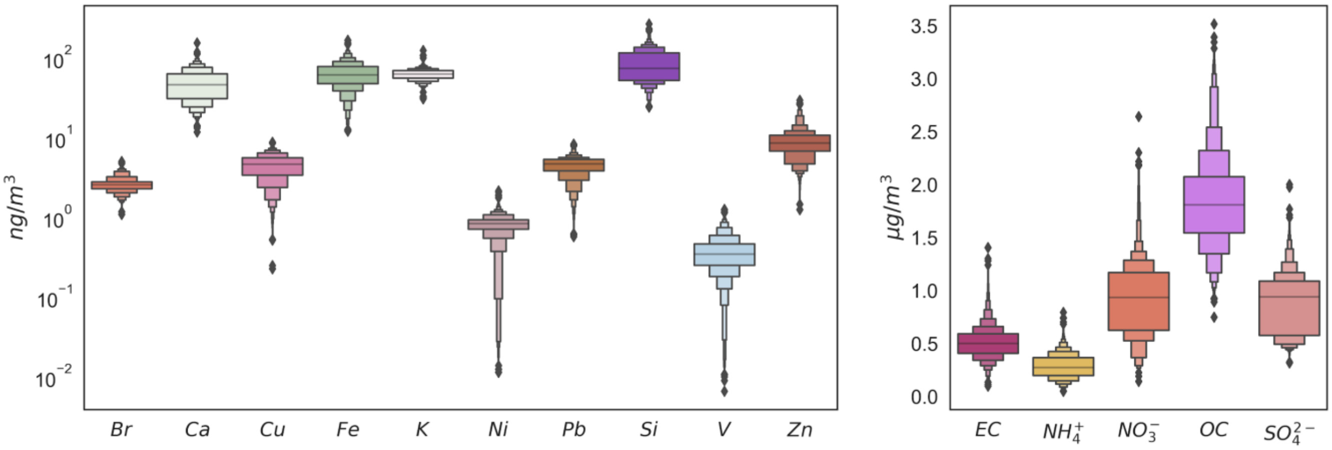


***Supplementary Figure 3. Mass concentrations of each $PM_{2.5}$ component.*** *Abbreviations*: bromine, Br; calcium, Ca; copper, Cu; elemental carbon, EC; iron, Fe; potassium, K; ammonium, $NH_4^+$; nickel, Ni; nitrate, $NO_3^+$; organic carbon, OC; lead, Pb; silicon, Si; sulfate, $SO_4^{2-}$; vanadium, V; zinc, Zn; fine particulate matter <2.5μm, $PM_{2.5}$.

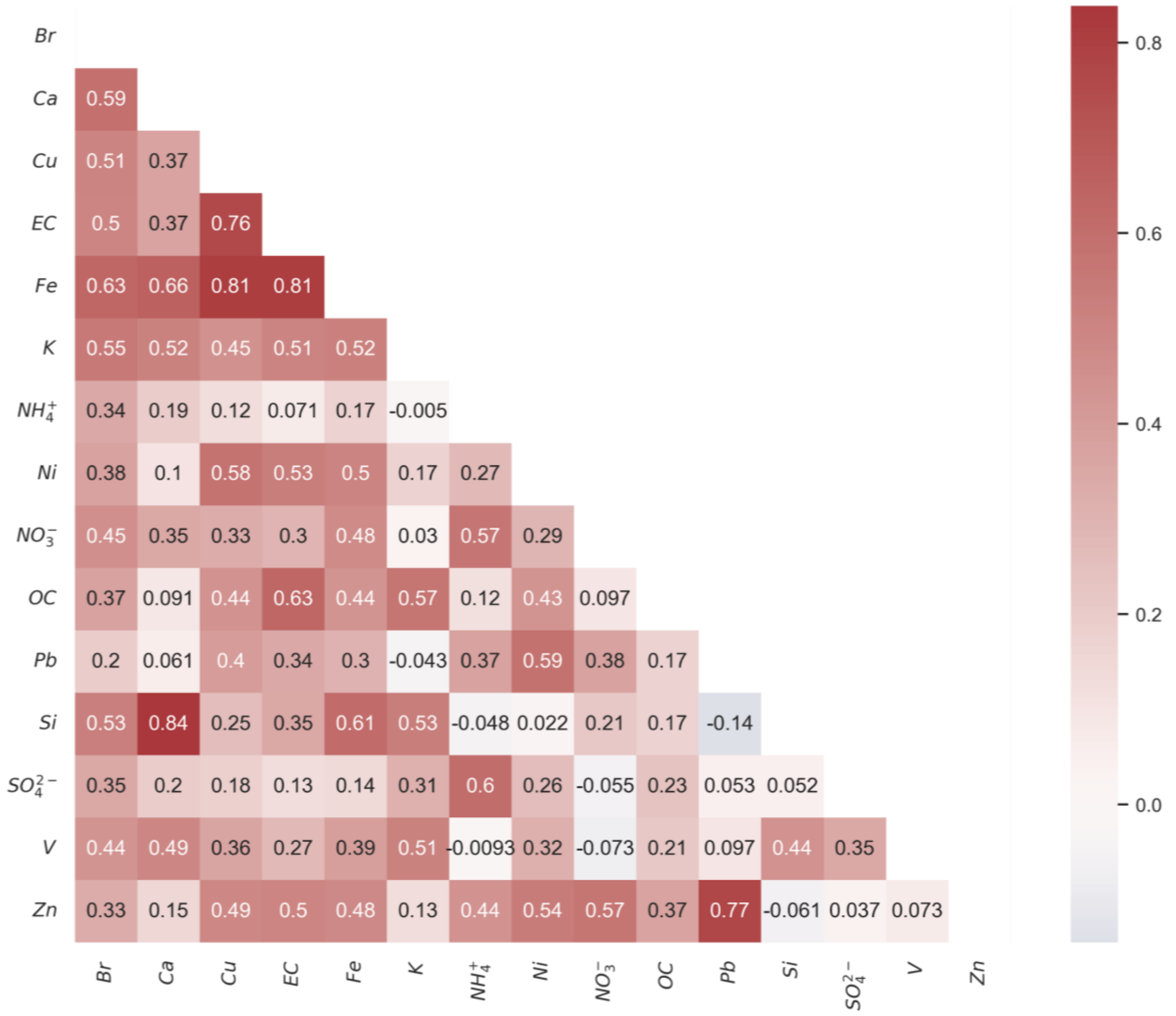


***Supplementary Figure 4. Correlations between components of $PM_{2.5}$.*** Spearman correlations between participant exposures to each source of $PM_{2.5}$ assessed here, derived from hybrid spatiotemporal modes (*4*, *5*) and assigned via participants' residential addresses. *Abbreviations*: bromine, Br; calcium, Ca; copper, Cu; elemental carbon, EC; iron, Fe; potassium, K; ammonium, $NH_4^+$; nickel, Ni; nitrate, $NO_3^-$; organic carbon, OC; lead, Pb; silicon, Si; sulfate, $SO_4^{2-}$; vanadium, V; zinc, Zn; fine particulate matter <2.5µm, $PM_{2.5}$.

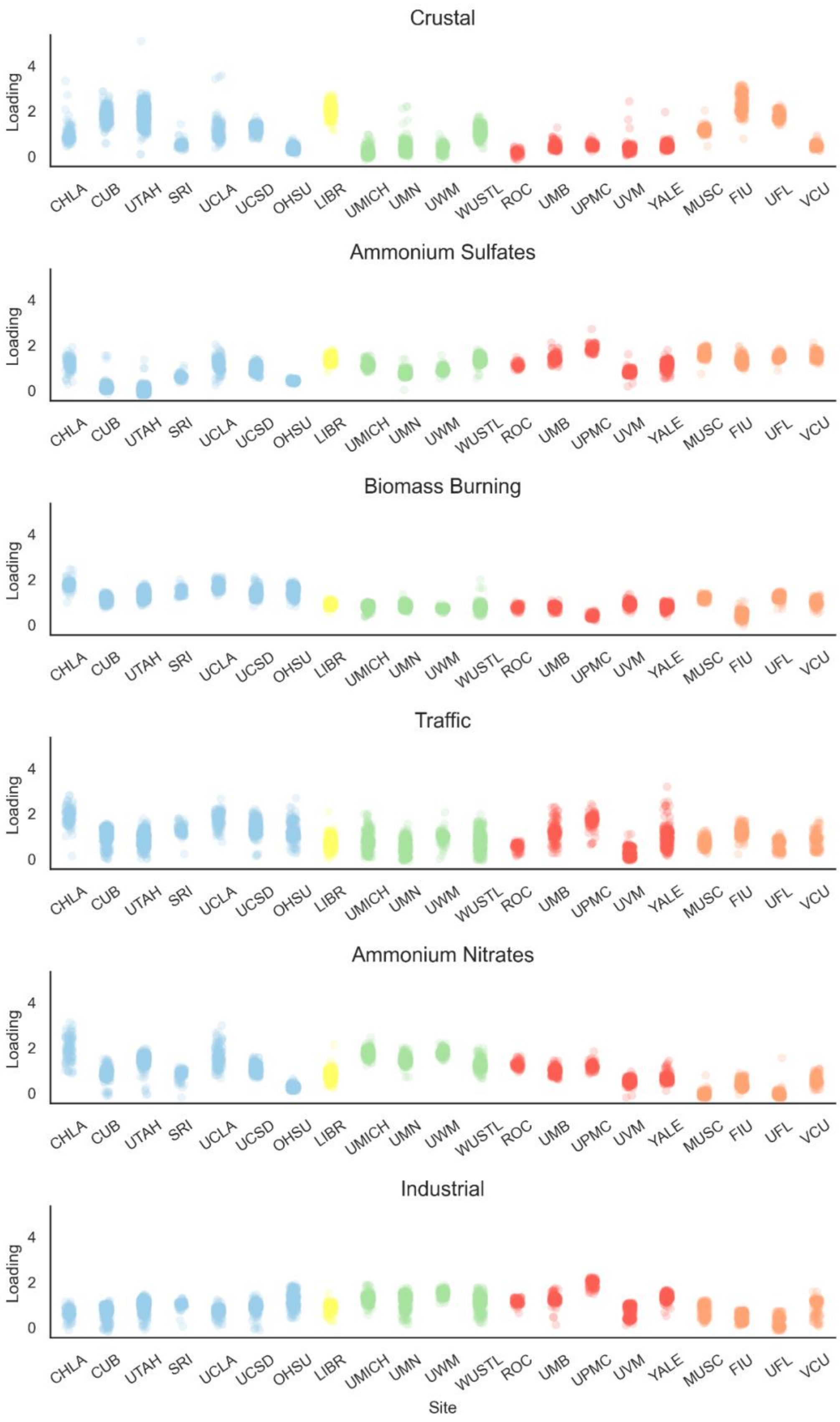
Crustal
Ammonium Sulfates
Biomass Burning
Traffic
Ammonium Nitrates
Industrial
Loading
0
2
4
CHLA
CUB
UTAH
SRI
UCLA
UCSD
OHSU
LIBR
UMICH
UMN
UWM
WUSTL
ROC
UMB
UPMC
UVM
YALE
MUSC
FIU
UFL
VCU
Site

***Supplementary Figure 5. Participant loadings per PMF-derived source of $PM_{2.5}$, by data collection site.*** Colors represent geographical regions across the United States. Blue = West: Children's Hospital Los Angeles (CHLA), University of Colorado Boulder (CUB), University of Utah (UTAH), SRI International (SRI), University of California Los Angeles (UCLA), University of California San Diego (UCSD), Oregon Health & Science University (OHSU); Yellow = Southwest: Laureate Institute for Brain Research (LIBR); Green = Midwest: University of Michigan (UMICH), University of Minnesota (UMN), University of Wisconsin-Milwaukee (UWM), Washington University in St. Louis (WUSTL); Red = Northeast: University of Rochester (ROC), University of Maryland at Baltimore (UMB), University of Pittsburgh (UPMC), University of Vermont (UVM), Yale University (YALE); Orange = Southeast: Medical University of South Carolina (MUSC), Florida International University (FIU), University of Florida (UFL), Virginia Commonwealth University (VCU).

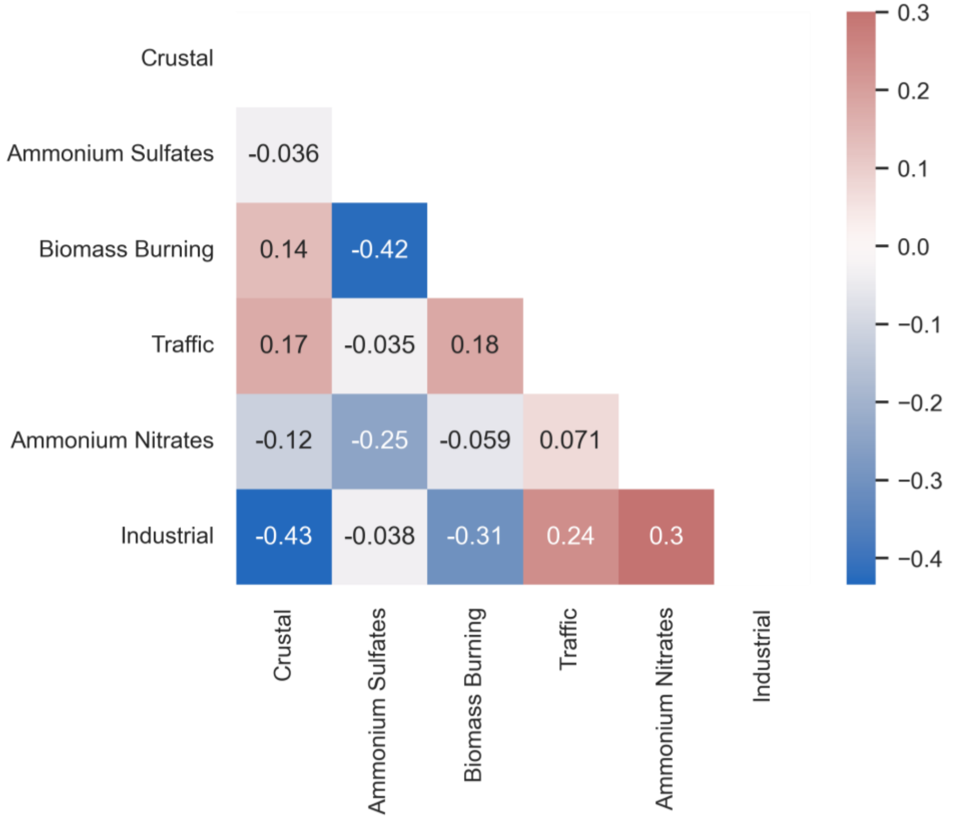


***Supplementary Figure 6. Correlations between loadings on each PMF-derived source of $PM_{2.5}$.*** Spearman correlations between participant loadings on each source of $PM_{2.5}$ as derived by positive matrix factorization (Sukumaran et al., under review).

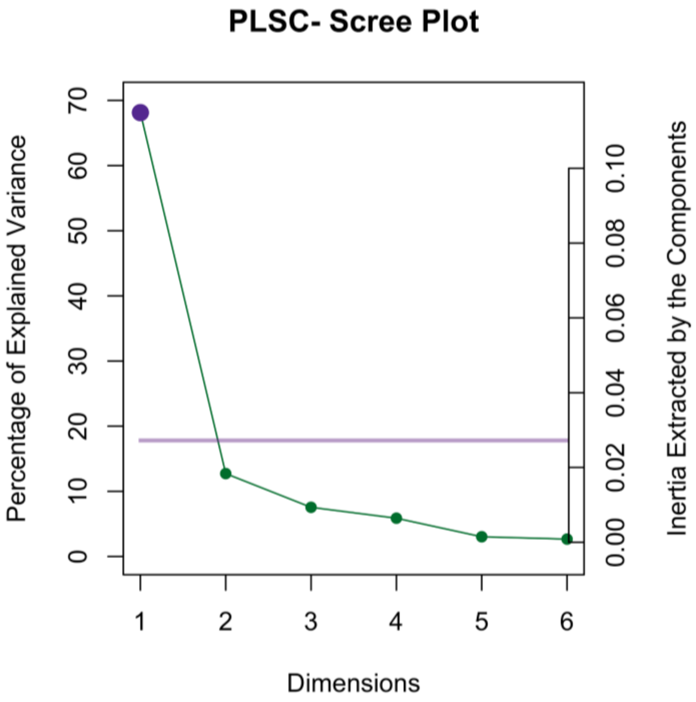


***Supplementary Figure 7.*** Scree plot showing shared variance in $PM_{2.5}$ sources and changes in anisotropic intracellular diffusion explained by each latent dimension. *Abbreviation*: partial least squares correlation, PLSC.

***Supplementary Table 5.*** Partial least squares correlation model statistics and eigenvalues.

| Model | Isotropic intracellular diffusion | | Anisotropic intracellular diffusion | |
|---|---|---|---|---|
| | Sources | Components | Sources | Components |
| *p*(Model) | 0.527 | 0.0001 | 0.0001 | 0.0001 |
| $\alpha_{FDR\text{-}corrected}$ | 0.0018 | 0.0007 | 0.0018 | 0.0007 |
| Proportion of variance, $\tau$ | | | | |
| 1 | 42% | ***35%*** | ***68%*** | ***74%*** |
| 2 | 21% | ***23%*** | 13% | ***9%*** |
| 3 | 16% | ***13%*** | 8% | 4% |
| 4 | 9% | 7% | 6% | 3% |
| 5 | 7% | 6% | 3% | 2% |
| 6 | 5% | 4% | 3% | 2% |
| 7 | – | 3% | – | 2% |
| 8 | – | 2% | – | 1% |
| 9 | – | 2% | – | 1% |
| 10 | – | 1% | – | 1% |
| 11 | – | 1% | – | 1% |
| 12 | – | 1% | – | 1% |
| 13 | – | 1% | – | 0% |
| 14 | – | 0% | – | 0% |
| 15 | – | 0% | – | 0% |

*Note.* Only latent dimensions of significant overall models are considered significant, even if eigenvalues suggested more significant latent dimensions. Component models have more potential latent factors than source models, as PLS limits the number of identifiable latent factors to the minimum number of variables between blocks (here: 68 brain regions vs. 6 sources/15 components). Type I error rate given for the number of effective comparisons (*6*) (*4*) between latent dimensions, at $\alpha < 0.01$.

*nentary Table 6*. Pollution source and brain region saliences, first latent dimension between pollution and anisotropic intracellular diffusion

| Variable | | Salience |
|---|---|---|
| Pollution source | | |
| Crustal | | -0.373 |
| Ammonium nitrates | | -0.184 |
| ***Biomass burning*** | | ***-0.763*** |
| Traffic | | -0.307 |
| Ammonium sulfates | | 0.367 |
| Industrial fuel burning | | -0.127 |
| ***Brain region*** | | |
| R | Postcentral Gyrus | -0.20 |
| L | Inferior Frontal Gyrus, Pars Opercularis | -0.20 |
| R | Precentral Gyrus | -0.19 |
| L | Postcentral Gyrus | -0.19 |
| L | Supramarginal Gyrus | -0.18 |
| R | Inferior Frontal Gyrus, Pars Opercularis | -0.18 |
| R | Superior Frontal Gyrus | -0.16 |
| L | Transverse Temporal Gyrus | -0.16 |
| L | Cingulate Gyrus, Rostral Anterior | -0.16 |
| R | Transverse Temporal Gyrus | -0.16 |
| L | Precentral Gyrus | -0.16 |
| L | Precuneus | -0.16 |
| R | Middle Frontal Gyrus, Rostral | -0.16 |
| L | Fusiform Gyrus | -0.15 |
| R | Supramarginal Gyrus | -0.15 |
| R | Superior Temporal Lobule | -0.15 |
| R | Insula | -0.15 |
| L | Cingulate Gyrus, Caudal Anterior | -0.15 |
| L | Orbitofrontal Gyrus , Lateral | -0.14 |
| L | Superior Frontal Gyrus | -0.13 |
| R | Middle Frontal Gyrus, Caudal | -0.13 |
| L | Middle Frontal Gyrus, Rostral | -0.13 |
| L | Cingulate Gyrus, Caudal Anterior | -0.13 |
| L | Inferior Parietal Lobule | -0.13 |
| R | Inferior Frontal Gyrus, Pars Triangularis | -0.13 |
| L | Middle Frontal Gyrus, Caudal | -0.12 |
| R | Orbitofrontal Gyrus, Lateral | -0.12 |
| L | Cuneus | -0.12 |
| R | Banks of Superior Temporal Sulcus | -0.12 |
| R | Inferior Parietal Lobule | -0.12 |
| R | Middle Temporal Gyrus | -0.12 |
| R | Precuneus | -0.12 |
| L | Inferior Frontal Gyrus, Pars Orbitalis | -0.12 |
| L | Inferior Frontal Gyrus, Pars Triangularis | -0.11 |
| L | Inferior Temporal Gyrus | -0.11 |
| R | Cuneus | -0.11 |

| | | |
|---|---|---|
| L | Banks of Superior Temporal Sulcus | -0.11 |
| L | Entorhinal Cortex | -0.11 |
| L | Superior Temporal Lobule | -0.11 |
| L | Parahippocampal Gyrus | -0.11 |
| R | Superior Parietal Lobule | -0.10 |

*Note.* All pollution sources and their saliences are noted. Bold values are significant at $p < 0.01$, bold and italicized values are significant at $p < 0.001$. Only significant brain regions' saliences are noted, all at $p < 0.01$. Abbreviations: L=left hemisphere; R=right hemisphere.

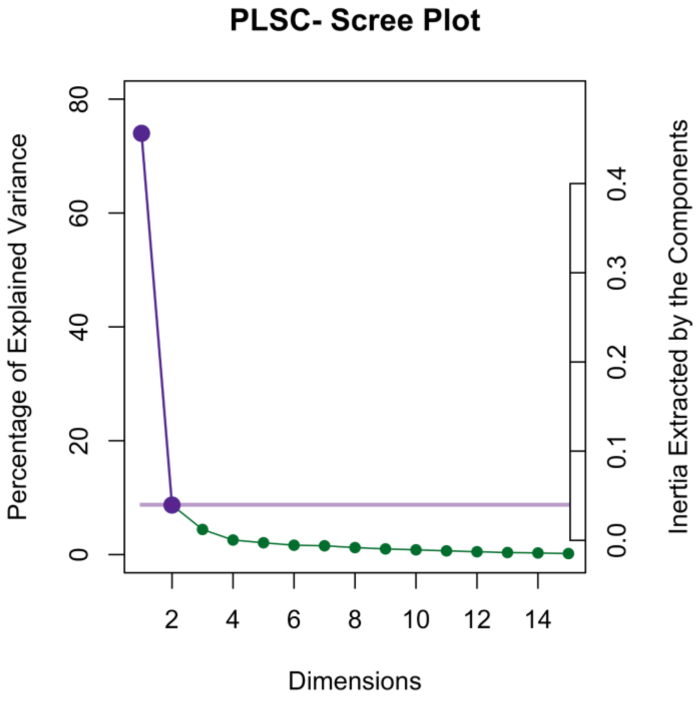


***Supplementary Figure 8***. Scree plot showing shared variance in PM components and changes in anisotropic intracellular diffusion explained by each latent dimension

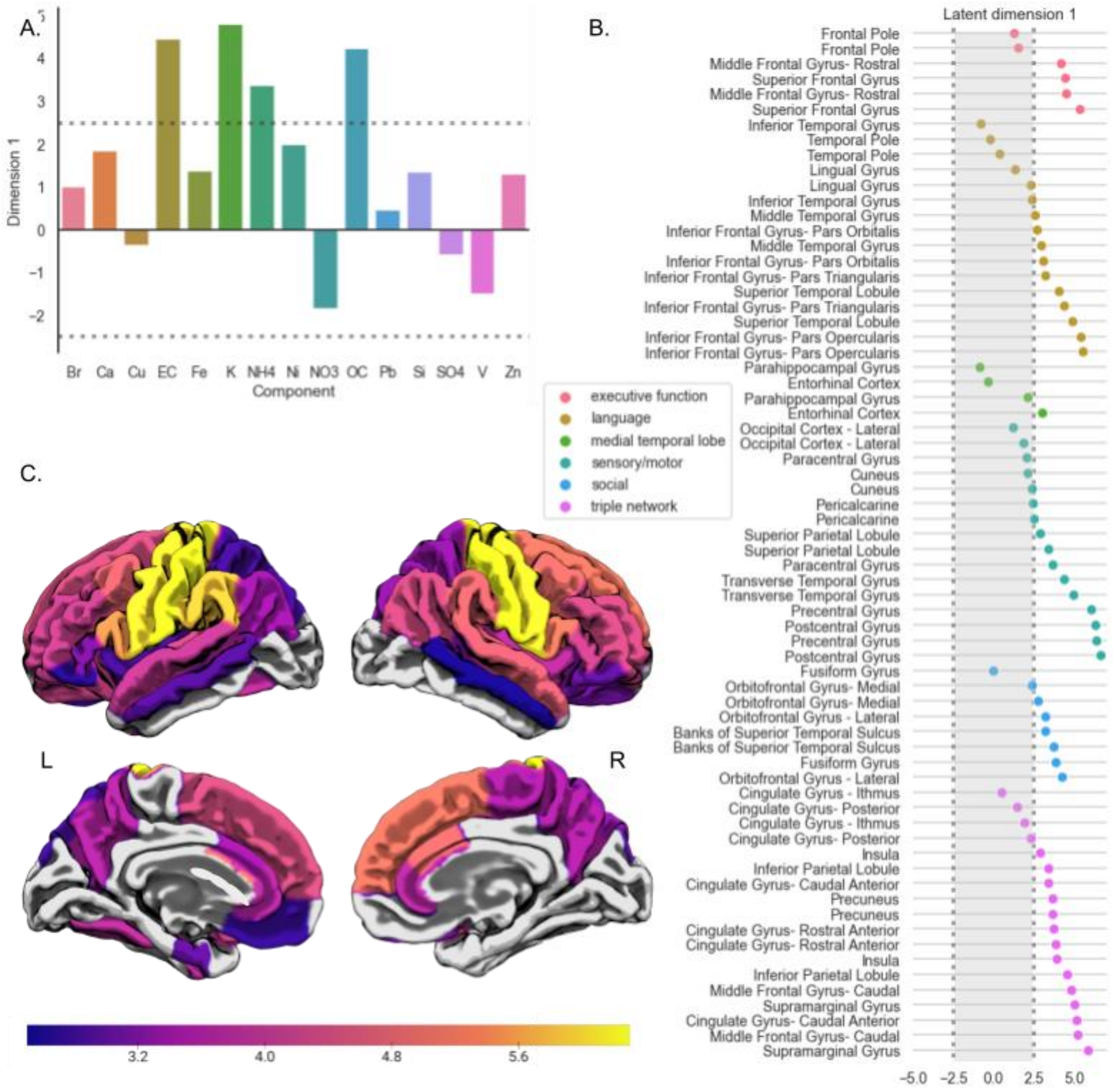
A.
Dimension 1
Component
Br
Ca
Cu
EC
Fe
K
NH4
Ni
NO3
OC
Pb
Si
SO4
V
Zn
B.
Latent dimension 1
executive function
language
medial temporal lobe
sensory/motor
social
triple network
Frontal Pole
Middle Frontal Gyrus- Rostral
Superior Frontal Gyrus
Inferior Temporal Gyrus
Temporal Pole
Lingual Gyrus
Middle Temporal Gyrus
Inferior Frontal Gyrus- Pars Orbitalis
Inferior Frontal Gyrus- Pars Triangularis
Superior Temporal Lobule
Inferior Frontal Gyrus- Pars Opercularis
Parahippocampal Gyrus
Entorhinal Cortex
Occipital Cortex - Lateral
Paracentral Gyrus
Cuneus
Pericalcarine
Superior Parietal Lobule
Transverse Temporal Gyrus
Precentral Gyrus
Postcentral Gyrus
Fusiform Gyrus
Orbitofrontal Gyrus- Medial
Orbitofrontal Gyrus - Lateral
Banks of Superior Temporal Sulcus
Cingulate Gyrus - Ithmus
Cingulate Gyrus- Posterior
Insula
Inferior Parietal Lobule
Cingulate Gyrus- Caudal Anterior
Precuneus
Cingulate Gyrus- Rostral Anterior
Middle Frontal Gyrus- Caudal
Supramarginal Gyrus
C.
L
R
3.2
4.0
4.8
5.6

***Supplementary Figure 9. First latent dimension of association between PM components and*** changes in anisotropic intracellular diffusion**.** A. Bootstrap ratios, equivalent to *z*-scores, indicate significant loadings (|*z*| > 2.5, dashed line) of air pollution components on the first latent dimension of brain-pollution associations. B. Bootstrap ratios indicate significant loadings (|*z*| > 2.5; outside the shaded box) of annualized change in anisotropic intracellular diffusion, across brain regions, on the first latent dimension of brain-pollution associations. Regional loading markers are color-coded based on broad functional categories of the functions of each brain region. C. The same regional loadings (i.e., bootstrap ratios, equivalent to *z*-scores) of significant changes in anisotropic intracellular diffusion, by region, on the first latent dimension of brain-pollution association, as shown in (B), plotted on a cortical surface template.

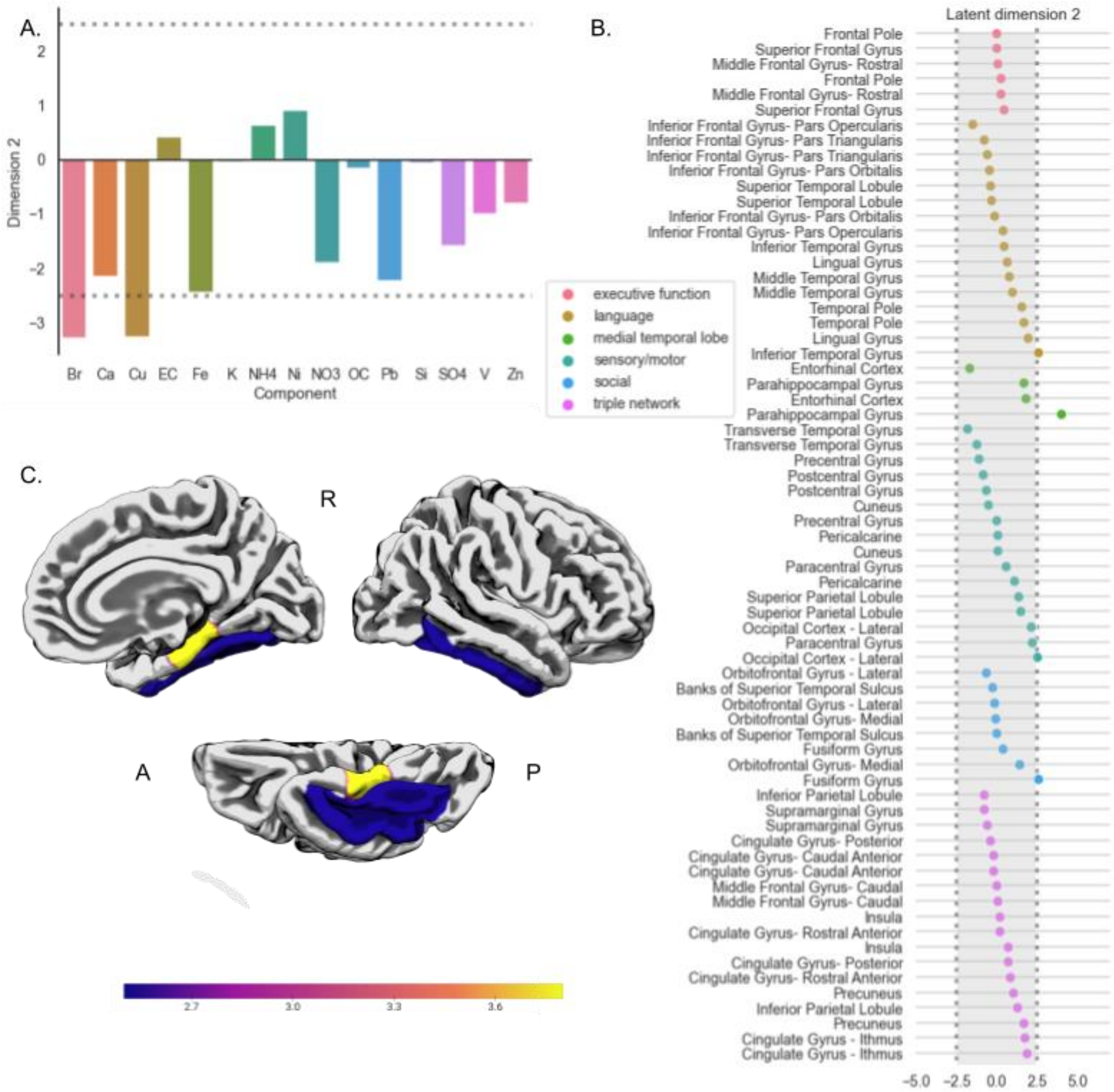


***Supplementary Figure 10. Second latent dimension of association between PM components and*** changes in anisotropic intracellular diffusion. A. Bootstrap ratios, equivalent to *z*-scores, indicate significant loadings (|*z*| > 2.5, dashed line) of air pollution components on the second latent dimension of brain-pollution associations. B. Bootstrap ratios indicate significant loadings (|*z*| > 2.5; outside the shaded box) of annualized change in anisotropic intracellular diffusion, across brain regions, on the second latent dimension of brain-pollution associations. Regional loading markers are color-coded based on broad functional categories of the functions of each brain region. C. The same regional loadings (i.e., bootstrap ratios, equivalent to *z*-scores) of significant changes in anisotropic intracellular diffusion, by region, on the second latent dimension of brain-pollution association, as shown in (B), plotted on a cortical surface template.

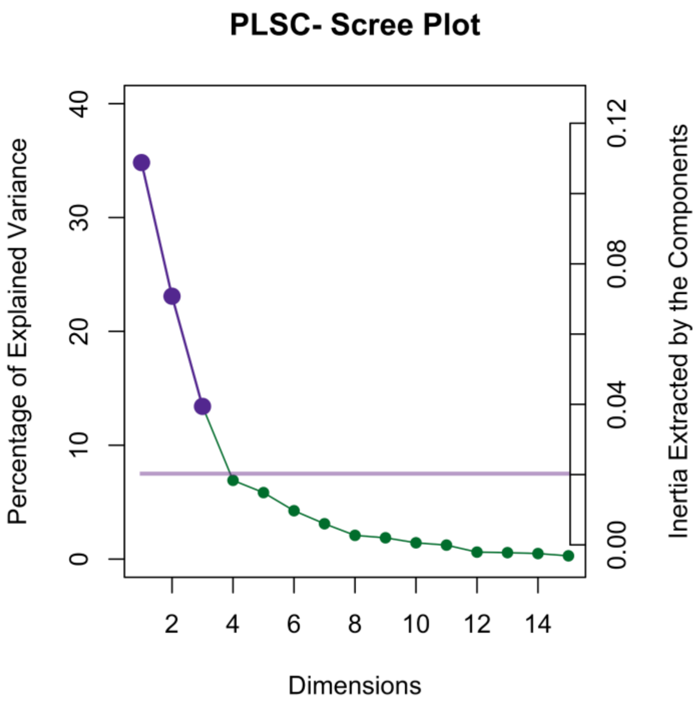


***Supplementary Figure 11***. Scree plot showing shared variance in PM components and changes in isotropic intracellular diffusion explained by each latent dimension

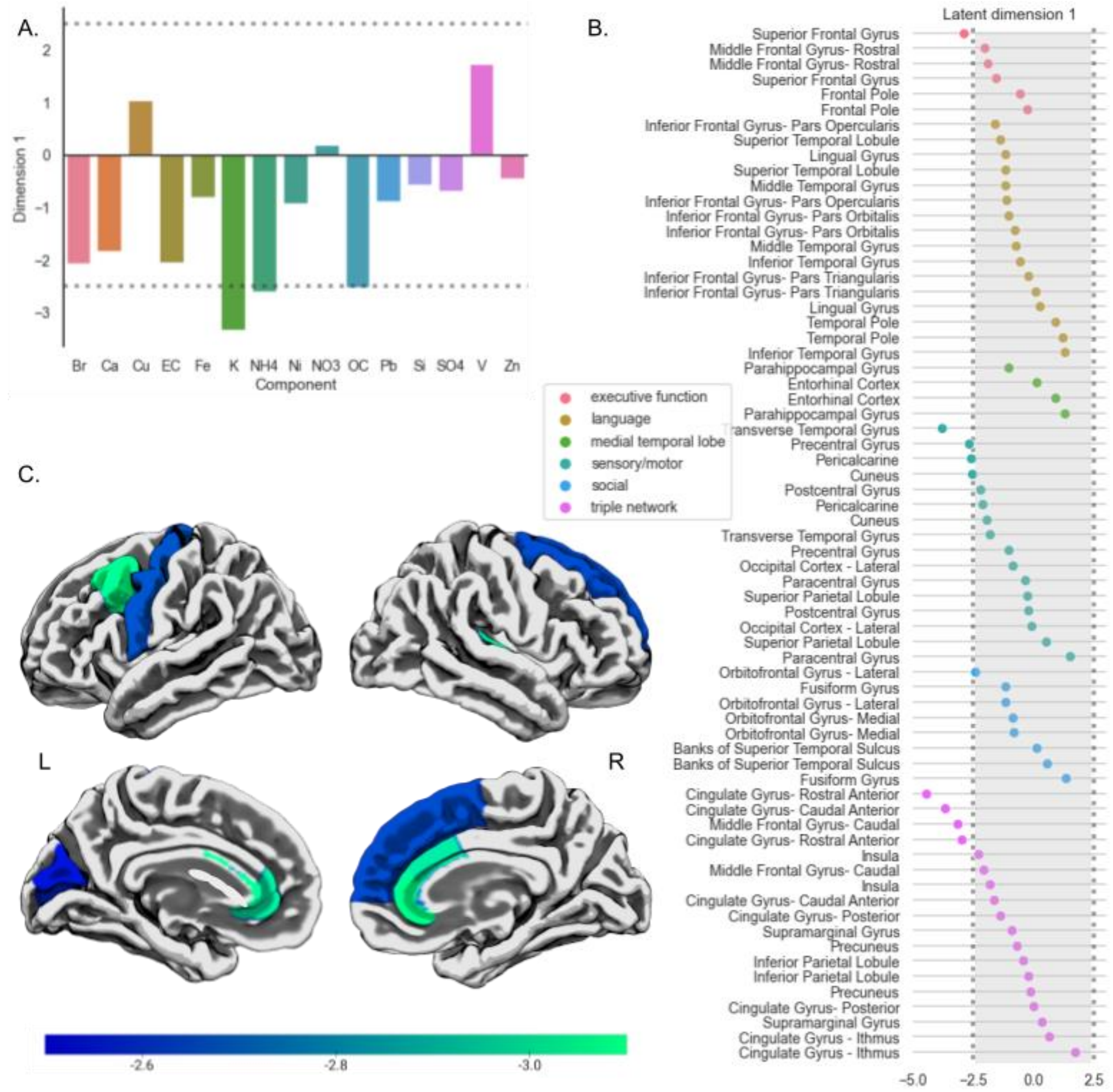


***Supplementary Figure 12. First latent dimension of association between PM components and changes in isotropic intracellular diffusion.*** A. Bootstrap ratios, equivalent to *z*-scores, indicate significant loadings (|*z*| > 2.5, dashed line) of air pollution components on the first latent dimension of brain-pollution associations. B. Bootstrap ratios indicate significant loadings (|*z*| > 2.5; outside the shaded box) of annualized change in isotropic intracellular diffusion, across brain regions, on the first latent dimension of brain-pollution associations. Regional loading markers are color-coded based on broad functional categories of the functions of each brain region. C. The same regional loadings (i.e., bootstrap ratios, equivalent to *z*-scores) of significant changes in isotropic intracellular diffusion, by region, on the first latent dimension of brain-pollution association, as shown in (B), plotted on a cortical surface template.

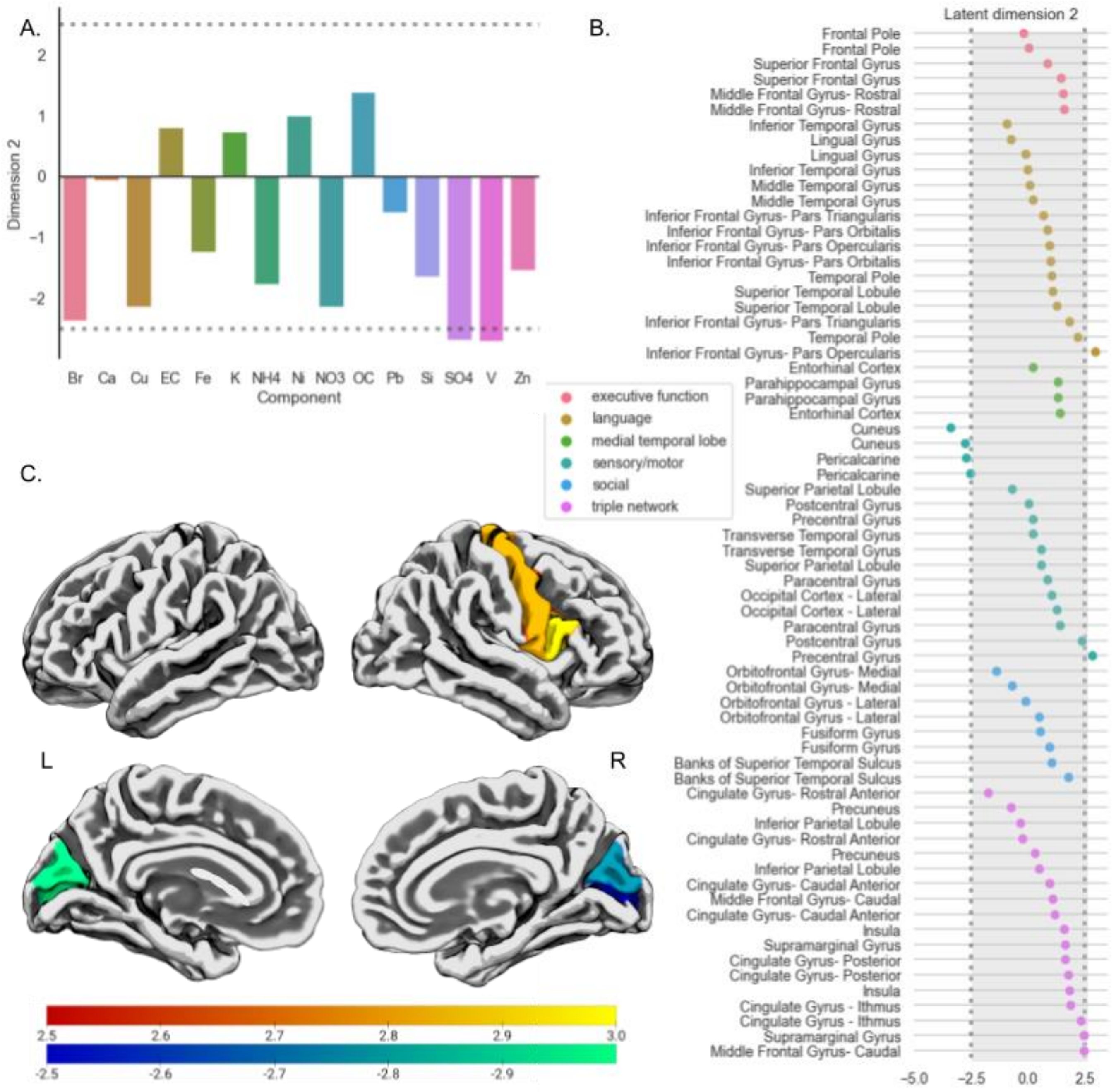
A.
B.
C.
Dimension 2
Component
Br
Ca
Cu
EC
Fe
K
NH4
Ni
NO3
OC
Pb
Si
SO4
V
Zn
executive function
language
medial temporal lobe
sensory/motor
social
triple network
L
R
Latent dimension 2
Frontal Pole
Frontal Pole
Superior Frontal Gyrus
Superior Frontal Gyrus
Middle Frontal Gyrus- Rostral
Middle Frontal Gyrus- Rostral
Inferior Temporal Gyrus
Lingual Gyrus
Lingual Gyrus
Inferior Temporal Gyrus
Middle Temporal Gyrus
Middle Temporal Gyrus
Inferior Frontal Gyrus- Pars Triangularis
Inferior Frontal Gyrus- Pars Orbitalis
Inferior Frontal Gyrus- Pars Opercularis
Inferior Frontal Gyrus- Pars Orbitalis
Temporal Pole
Superior Temporal Lobule
Superior Temporal Lobule
Inferior Frontal Gyrus- Pars Triangularis
Temporal Pole
Inferior Frontal Gyrus- Pars Opercularis
Entorhinal Cortex
Parahippocampal Gyrus
Parahippocampal Gyrus
Entorhinal Cortex
Cuneus
Cuneus
Pericalcarine
Pericalcarine
Superior Parietal Lobule
Postcentral Gyrus
Precentral Gyrus
Transverse Temporal Gyrus
Transverse Temporal Gyrus
Superior Parietal Lobule
Paracentral Gyrus
Occipital Cortex - Lateral
Occipital Cortex - Lateral
Paracentral Gyrus
Postcentral Gyrus
Precentral Gyrus
Orbitofrontal Gyrus- Medial
Orbitofrontal Gyrus- Medial
Orbitofrontal Gyrus - Lateral
Orbitofrontal Gyrus - Lateral
Fusiform Gyrus
Fusiform Gyrus
Banks of Superior Temporal Sulcus
Banks of Superior Temporal Sulcus
Cingulate Gyrus- Rostral Anterior
Precuneus
Inferior Parietal Lobule
Cingulate Gyrus- Rostral Anterior
Precuneus
Inferior Parietal Lobule
Cingulate Gyrus- Caudal Anterior
Middle Frontal Gyrus- Caudal
Cingulate Gyrus- Caudal Anterior
Insula
Supramarginal Gyrus
Cingulate Gyrus- Posterior
Cingulate Gyrus- Posterior
Insula
Cingulate Gyrus - Ithmus
Cingulate Gyrus - Ithmus
Supramarginal Gyrus
Middle Frontal Gyrus- Caudal

***Supplementary Figure 13. Second latent dimension of association between PM components and changes in isotropic intracellular diffusion.*** A. Bootstrap ratios, equivalent to $z$-scores, indicate significant loadings ($|z| > 2.5$, dashed line) of air pollution components on the second latent dimension of brain-pollution associations. B. Bootstrap ratios indicate significant loadings ($|z| > 2.5$; outside the shaded box) of annualized change in isotropic intracellular diffusion, across brain regions, on the second latent dimension of brain-pollution associations. Regional loading markers are color-coded based on broad functional categories of the functions of each brain region. C. The same regional loadings (i.e., bootstrap ratios, equivalent to $z$-scores) of significant changes in isotropic intracellular diffusion, by region, on the second latent dimension of brain-pollution association, as shown in (B), plotted on a cortical surface template.

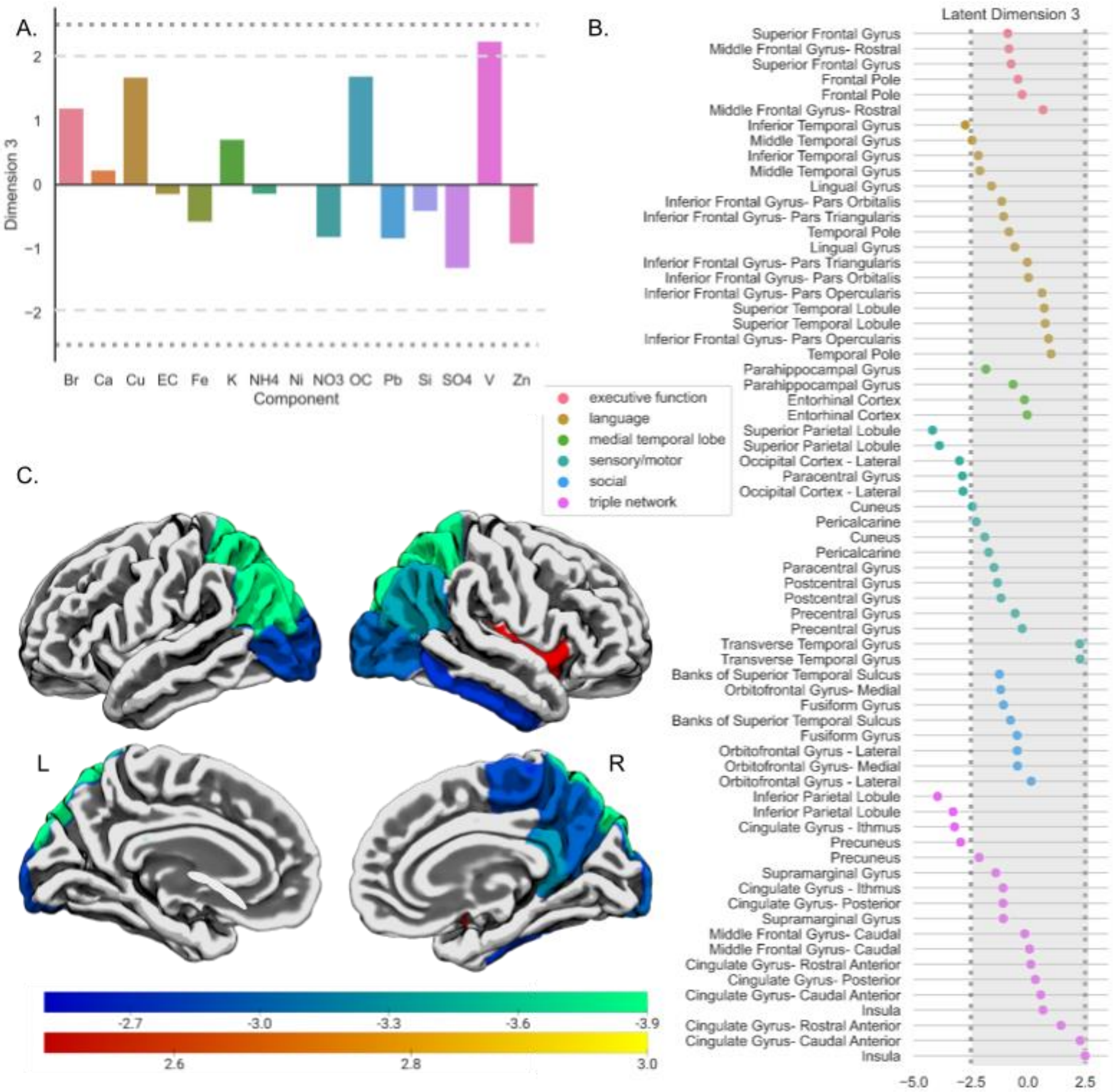


***Supplementary Figure 14**. Third latent dimension of association between PM components and changes in isotropic intracellular diffusion.* A. Bootstrap ratios, equivalent to *z*-scores, indicate significant loadings (|*z*| > 2.5, dashed line) of air pollution components on the third latent dimension of brain-pollution associations. B. Bootstrap ratios indicate significant loadings (|*z*| > 2.5; outside the shaded box) of annualized change in isotropic intracellular diffusion, across brain regions, on the third latent dimension of brain-pollution associations. Regional loading markers are color-coded based on broad functional categories of the functions of each brain region. C. The same regional loadings (i.e., bootstrap ratios, equivalent to *z*-scores) of significant changes in isotropic intracellular diffusion, by region, on the third latent dimension of brain-pollution association, as shown in (B), plotted on a cortical surface template.

#### References

1. I. Shrier, R. W. Platt, Reducing bias through directed acyclic graphs. *BMC Med. Res. Methodol.* **8**, 70 (2008).
2. J. Textor, B. van der Zander, M. S. Gilthorpe, M. Liskiewicz, G. T. Ellison, Robust causal inference using directed acyclic graphs: the R package "dagitty." *Int. J. Epidemiol.* **45**, 1887–1894 (2016).
3. H. Amini, M. Danesh-Yazdi, Q. Di, W. Requia, Y. Wei, Y. AbuAwad, L. Shi, M. Franklin, C. M. Kang, M. J. Wolfson, Annual mean PM2. 5 components (EC, NH4, NO3, OC, SO4) 50m urban and 1km non-urban area grids for contiguous US, 2000-2019 v1. *Palisades N. Y. NASA Socioecon. Data Appl. Cent. SEDAC* (2023).
4. Q. Di, H. Amini, L. Shi, I. Kloog, R. Silvern, J. Kelly, M. B. Sabath, C. Choirat, P. Koutrakis, A. Lyapustin, An ensemble-based model of PM2. 5 concentration across the contiguous United States with high spatiotemporal resolution. *Environ. Int.* **130**, 104909 (2019).
5. Q. Di, H. Amini, L. Shi, I. Kloog, R. Silvern, J. Kelly, M. B. Sabath, C. Choirat, P. Koutrakis, A. Lyapustin, Y. Wang, L. J. Mickley, J. Schwartz, Assessing NO2 Concentration and Model Uncertainty with High Spatiotemporal Resolution across the Contiguous United States Using Ensemble Model Averaging. *Environ. Sci. Technol.* **54**, 1372–1384 (2020).
6. J. Li, L. Ji, Adjusting multiple testing in multilocus analyses using the eigenvalues of a correlation matrix. *Heredity* **95**, 221–227 (2005).
